# Making more with forest inventory data: Toward a scalable, dynamical model of forest change

**DOI:** 10.1101/2024.07.22.604669

**Authors:** Malcolm Itter, Andrew O. Finley

## Abstract

Models of forest dynamics are an important tool to understand and predict forest responses to global change. Despite recent model development, predictions of forest dynamics under global change remain highly variable reflecting uncertainty in future conditions, forest demographic processes, and the data used to parameterize and validate models. Quantifying this uncertainty and accounting for it when making adaptive management decisions is critical to our ability to conserve forest ecosystems in the face of rapidly changing conditions. Dynamical spatio-temporal models (DSTMs) are a particularly powerful tool in this setting given they quantify uncertainty associated with process-based models of forest demography, the parameters upon which those models depend, and the forest data used to inform them. Further, DSTMs propagate this uncertainty to predictions of forest dynamics allowing for its formal integration within adaptive management decision frameworks. A major challenge to the application of DSTMs in applied forest ecology has been the lack of a scalable, theoretical model of forest dynamics that generates predictions at the stand level—the scale at which management decisions are made. We address this challenge by integrating a matrix projection model motivated by the McKendrick-von Foerster partial differential equation for size-structured population dynamics within a Bayesian hierarchical DSTM informed by forest inventory data. The model provides probabilistic predictions of species-specific demographic rates and changes in the size-species distribution over time. The model is applied to predict long-term dynamics (60+ years) within the Penobscot Experimental Forest in Maine, USA, accounting for uncertainty in inventory observations, process-based predictions, and model parameters for nine Acadian Forest species. We find that variability in inventory observations associated with heterogeneous stand conditions drives uncertainty in predictions of forest dynamics. We conclude with a discussion of how the initial DSTM can be refined and extended to better represent forest dynamics under global change and inform adaptive management.

## Introduction

Models of forest dynamics (FDMs) have a long history in applied forest ecology providing predictions of forest demography, composition, structure, and function over time and space (Bugmann, 2001). They are applied in a range of settings including to assess alternative management strategies (under the broad heading of growth and yield modeling; Pretzsch, 2009), to test mechanisms of species co-existence (Falster et al., 2017), and to predict complex biogeochemical processes at large spatial scales as part of dynamic global vegetation models (Fisher et al., 2018). Contemporary FDMs are also an important adaptive management tool, informing actions to promote ecosystem resilience and maintain valuable ecosystem functions under rapidly changing conditions (Trasobares et al., 2016; Radke et al., 2020).

Global change is already contributing to fundamental shifts in historic forest dynamics making it difficult to predict the future state of forest ecosystems (McDowell et al., 2020). While contemporary FDMs are generally consistent in their approximation of historic forest dynamics, predictions under future climate and disturbance scenarios vary widely among models (Fisher et al., 2010; Bugmann et al., 2019). This variation is driven by uncertainty in future conditions including altered climate and disturbance regimes, imperfect and variable representation of the demographic processes underlying forest dynamics, and noisy observations of forests under novel conditions (Bugmann and Seidl, 2022). From a management perspective, this increased uncertainty means that historic patterns of forest dynamics, which serve as the basis for sustainable management approaches, are not necessarily good indicators of the future state of forest ecosystems.

Accounting for the uncertainty in future forest dynamics is critical to making effective adaptive management decisions to conserve forest ecosystems and the myriad services they provide (Messier et al., 2016). While there has been rapid advancement of FDMs in recent decades to improve the accuracy of predictions of forest dynamics under global change, accounting for the uncertainty in these predictions remains challenging (Bugmann and Seidl, 2022). In particular, the size and complexity of modern FDMs makes them difficult to integrate within probabilistic frameworks aimed at quantifying different sources of uncertainty and propagating it to predictions of forest dynamics (Dietze et al., 2018). Thus, there is a need for a probabilistic FDM that generates predictions of forest dynamics with explicit estimates of uncertainty that can be used to inform adaptive forest management decisions (Yousefpour et al., 2012).

Here, we develop a dynamical statistical model that provides probabilistic predictions of future forest states at the spatial scale at which management decisions are made in terms directly implementable by forest managers. The outputs of the model can be used within adaptive management decision frameworks that support optimal decision making in the face of uncertain future conditions (Yousefpour et al., 2012; Radke et al., 2020). To provide context and motivation for our modeling approach, we first review methods to quantify uncertainty within FDMs, detail the fundamental differences between these methods and a dynamical statistical model, and describe the importance of prediction scale to support adaptive management decisions.

### Uncertainty in predictions of forest dynamics

Uncertainty in predictions of forest dynamics stems from multiple sources including the data used to parameterize and drive FDMs (data uncertainty), the functions used to approximate the ecological processes underlying forest dynamics (process uncertainty), unknown model parameters that must be estimated in order to generate predictions including those representing the initial conditions of a forest ecosystem (parameter uncertainty Dietze, 2017; Van Oijen, 2017). Data uncertainty reflects the fact that the state of forest ecosystems and the factors that drive forest dynamics (e.g., disturbance, climate) are never observed perfectly. Process uncertainty reflects incomplete understanding of the complex processes underlying forest dynamics and accounts for potential model misspecification. Uncertainty in model parameters reflects their imperfect estimation based on noisy data. Lastly, initial condition uncertainty reflects the fact that the starting point for predictions of forest dynamics is often unknown and must be estimated either from data or an existing model (Raiho et al., 2020).

Ideally, FDMs would provide predictions of forest dynamics that reflect all of the uncertainty sources described above. In practice, however, the structure of FDMs including multiple non-linear demographic functions with potentially correlated parameters along with non-Gaussian observations make it challenging to quantify and propagate uncertainty to predictions of forest dynamics (Wilson et al., 2019). Given the large number of parameters in FDMs, the specific data needed to inform them, and the potential for unmodeled correlation among parameters to limit their identifiability, FDMs are often parameterized independently for different demographic processes (growth, mortality, regeneration) and species (Purves et al., 2008). This independent approach to FDM parameterization limits uncertainty propagation given the challenges of incorporating estimates of parameter uncertainty from separate model fits into joint predictions of forest dynamics (Myllymäki et al., 2024). As a result, FDMs commonly provide point predictions—a single predicted value representing the best guess of the future of a forest ecosystem (Fisher et al., 2010; Bugmann et al., 2019). A simple example of point prediction is to fix FDM parameters at maximum likelihood estimates and generate predictions approximating the mean of the future state of the forest. Point predictions do not reflect uncertainty in the future state of forest ecosystems as there is no variability in predicted values (Gneiting, 2011).

The alternative to point prediction is probabilistic prediction, under which a probability distribution is generated for each predicted value reflecting uncertainty in future forest states (Gneiting and Katzfuss, 2014). The generated probability distribution provides a measure of variability in predicted values (Dietze, 2017). Intermediate between point and probabilistic prediction are FDMs that incorporate variability in demographic rates at the individual tree or forest patch scale as a measure of demographic stochasticity or local environmental heterogeneity (Fisher et al., 2018). Note that while the inclusion of demographic stochasticity results in variability in predicted forest demography (most commonly growth), it is not equivalent to probabilistic prediction under which predicted forest ecosystem states follow a formal probability distribution reflecting all sources of uncertainty included in the model.

In general, it is not possible to determine the probability distribution of predicted forest states directly given the complex structure of FDMs. Instead, probabilistic predictions of forest dynamics are generated numerically through techniques such as bootstrapping and Monte Carlo simulation (Van Oijen, 2017). These techniques are computationally demanding requiring estimates of uncertainty for one or more of the sources described above (data, process, parameters) and, as a result, are less commonly applied. When probabilistic prediction is applied, uncertainty quantification is frequently limited to parameter uncertainty alone given it is more easily quantified than other uncertainty types. To account for parameter uncertainty, FDM parameters are sampled conditional on respective variance estimates and used to generate an ensemble of predicted future states reflecting uncertainty in underlying model parameters (Fisher et al., 2010).

Accounting for parameter uncertainty is more straightforward if forest dynamics are modeled under a Bayesian approach given there exists a posterior predictive distribution that integrates over the uncertainty in model parameters propagating it to predictions of forest states (Hartig et al., 2012). There are few examples of Bayesian FDMs under which all model parameters are estimated jointly allowing for probabilistic prediction reflecting parameter uncertainty (Purves et al., 2008; Wilson et al., 2019). Instead, demographic sub-models are often fit independently under Bayesian approaches, again necessitating an ensemble forecast to account for uncertainty in model parameters (Myllymäki et al., 2024). In some cases, uncertainty in future drivers of forest dynamics (e.g., climate, disturbance) are incorporated into probabilistic predictions (Spadavecchia et al., 2011). This is particularly challenging as it requires a probabilistic model for future scenarios. In lieu of such a probabilistic model, forest dynamics are sometimes predicted under a set of scenarios reflecting a non-probabilistic range of future conditions (Radke et al., 2020).

Other approaches to quantify uncertainty in predictions of forest dynamics include model inter-comparisons under which a selection of FDMs are used to make predictions of forest ecosystem change given a shared initial condition (Bugmann et al., 2019; Díaz-Yáñez et al., 2024). These inter-comparisons provide a measure of process uncertainty in FDMs, specifically the variable representation of the processes underlying forest dynamics. However, without performing some form of model averaging, these approaches do not allow for the propagation of process uncertainty to predictions. Further, to the extent that inter-comparisons utilize point rather than probabilistic predictions to compare FDMs, they do not account for the varying levels of uncertainty among models.

### Dynamical statistical models

An alternative approach to quantify and propagate uncertainty in predictions of forest dynamics is to apply a dynamical statistical model. These models pair a process model driving changes in a system over time and/or space with one or more data models that account for uncertainty in observations within a broader probabilistic framework. In the current context, an FDM represents the process model providing predictions of the state of forest ecosystems over time and space. These predictions are subject to an explicit process error term that reflects uncertainty in the mathematical representation of forest dynamics. Predictions are informed by observations of the state of forest ecosystems at select times and locations. These observations are imperfect and variable, and are subject to an explicit observation error term. Conceptually, forest observations are generated conditional on the latent (true, unobserved) state of the forest ecosystem plus some observation error. Hereafter, we refer to these models as dynamical spatio-temporal models (DSTMs) noting that in some contexts they are also referred to as state-space models (Wikle and Hooten, 2010). When implemented as part of a broader Bayesian hierarchical framework, DSTMs incorporate parameter models allowing for the quantification of parameter uncertainty in addition to the process and data uncertainty already described. In the context of modeling forest dynamics, Bayesian DSTMs generate probabilistic predictions of forest states that reflect uncertainty in the data, process, and parameters.

Despite the capacity of Bayesian DSTMs to quantify uncertainty in predictions of dynamic ecological processes (Laubmeier et al., 2020), there has been limited application of such models to predict forest dynamics over time and space. Relevant examples include models to predict individual tree demography (Clark et al., 2010, 2011), and to predict changes in the size distribution of forest populations over time using an integral projection process model (Ghosh et al., 2012). An important factor contributing to the limited application of Bayesian DSTMs to model forest dynamics is the scale and complexity of modern FDMs. As previously noted, these models are computationally demanding, utilizing a number of non-linear functions to approximate demographic processes often operating at the scale of individual trees. Integrating these complex models into Markov chain Monte Carlo (MCMC) routines commonly used to fit Bayesian DSTMs is generally intractable particularly over large spatio-temporal domains. Recent research has focused on the use of statistical emulators to reduce this computational challenge while still allowing for improved uncertainty quantification (Fer et al., 2018; Raiho et al., 2021).

An alternative approach to overcome the computational challenges of modeling complex spatio-temporal processes is to utilize low-dimensional model parameterizations motivated by process-based models within a broader Bayesian hierarchical framework (Wikle and Hooten, 2010). This approach relates to seminal applications of DSTMs in ecology (Wikle, 2003; Clark and Bjørnstad, 2004) and relies on conditioning facilitated by a hierarchical model structure to simplify complex ecological processes into component parts that are easier to model.

### The importance of scale

The scale at which forest dynamics are modeled is a key consideration in developing a computationally efficient DSTM. While individual-based approaches are common in current FDMs (Shugart et al., 2018), they greatly increase model dimensionality and make it difficult to scale models across multiple locations. The alternative is to approximate forest dynamics using stand-scale models, which are more computationally feasible to integrate within DSTMs given they generate population-level predictions (e.g., trees per hectare, basal area per hectare). Here, a forest stand is defined as a forest area approximately homogeneous in its species composition, size, structure, density, site quality, and management history (Ashton and Kelty, 2018).

Stand-scale models are also preferred if the goal is to use predictions of forest dynamics to inform adaptive management. The stand is the fundamental management unit and, despite its operational rather than ecological origins, remains the scale at which management decisions are made (O’Hara and Nagel, 2013). The stand is also the scale at which management for forest resilience and adaptation to global change are most often considered (Seidl et al., 2013). Aligning model predictions with stand-scale adaptive management outcomes is important both to inform management decisions (accounting for the uncertainty in predictions), but also to maximize the potential of new information gained through adaptive management experiments to reduce uncertainty in predictions (Dietze et al., 2018).

### Study overview

We develop a Bayesian hierarchical DSTM to model forest dynamics. The goal of the model is to provide probabilistic predictions of future forest states that can be used to make adaptive management decisions accounting for uncertainty in forest dynamics. To this end, we defined several objectives that were used to guide the structure, data, and implementation of the dynamical model: (1) the model should formally quantify data, process, and parameter (including parameters defining initial conditions) uncertainty and propagate this uncertainty to predictions of forest states; (2) forest dynamics should be modeled at a stand scale given its operational connection to adaptive management; (3) the model should be constructed using forest inventory data given its fundamental role in shaping management decisions (Burkhart et al., 2018); (4) the model should be scalable in the sense that it represent complex forest dynamics based on few parameters allowing the model to eventually be scaled up to predict the future of forest ecosystems across a region.

The initial model is defined to predict stand-scale forest dynamics over time informed by periodic inventory data. We apply the model to predict the evolution of stand conditions over 60 years within the Penobscot Experimental Forest. The initial model represents a proof-of-concept of uncertainty quantification, data integration, and probabilistic prediction in models of forest dynamics. We describe refinements and extensions to the initial dynamical model in the Discussion.

## Materials and methods

### Penobscot Experimental Forest inventory data

We develop our dynamical model using long-term inventory data from the Penobscot Experimental Forest (PEF). The PEF is a 1600 ha experimental forest north of Bangor, Maine (44^*°*^52’N, 68^*°*^38’ W) jointly managed by the USDA Forest Service and the University of Maine to support long-term applied forest ecology research (Brissette and Kenefic, 2014). The forest is located within the Acadian Forest, a transition zone between the eastern temperate forest to the south and the boreal forest to the north. The PEF is characterized by a cold, humid climate with a mean annual temperature of 6.8^*°*^C and mean annual precipitation of greater than 1000 mm with much falling as snow.

A long-term silviculture experiment was established in the PEF between 1952-1957 with replicated treatments initiated across 20 management units (Brissette and Kenefic, 2014). Permanent sample plots were installed prior to silvicultural treatments in the 1950s with plots remeasured every 5-10 years through present. Species identity and diameter at breast height (DBH) are recorded for all trees within sample plots using a nested, fixed area design with 0.08 ha plots used to measure large diameter trees (*>* 11.4 cm) and either 0.02 ha or 0.008 ha nested subplots used to measure small diameter trees (1.3 − 11.4 cm). Tree counts in 2.5 cm DBH classes were recorded in each plot prior to 1974 with continuous DBH measured to the nearest 0.3 cm in subsequent years.

We develop our dynamical model using previously published inventory data (Kenefic et al., 2015) collected within a 2.4 ha control unit in the PEF long-term silviculture experiment: MU32B (Fig. S1; hereafter referred to as the modeled stand). No silvicultural treatments were applied within this stand nor were there any major disturbance events during the model period. The stand is even-aged and characterized as being within the late stem exclusion to understory reinitiation phases of development over the model period (Brissette and Kenefic, 2014). The stand comprises a mixture of nine Acadian forest species including balsam fir (*Abies balsamea* (L.) Mill.), eastern hemlock (*Tsuga canadensis* (L.) Carr.), eastern white pine (*Pinus strobus* L.), northern white-cedar (*Thuja occidentalis* L.), paper birch (*Betula papyrifera* Marsh.), red maple (*Acer rubrum* L.), and red (*Picea rubens* Sarg.), black (*P. mariana* Mill.), and white spruce (*P. glauca* (Mo.) A.V.). The stand was inventoried 11 times between 1954-2009 (roughly every five years) across three (1954-1989) or ten (1993-2009) inventory plots (see Table S1 for inventory years and plot details).

### Dynamical model of forest change

We apply a cohort-based approach to model stand-scale forest dynamics. Cohort-based approaches connect to theoretical models of structured population dynamics (where structure may refer to species, size, and/or traits) and have been shown to well approximate complex forest dynamics (Strigul et al., 2008). Under qualifying assumptions, cohort-based models of forest dynamics can be expressed as the McKendrick-von Foerster partial differential equation (hereafter, “MvF PDE”; De Roos, 1997). The MvF PDE models changes in the density of individuals of a given species and size (and/or trait value) over time as

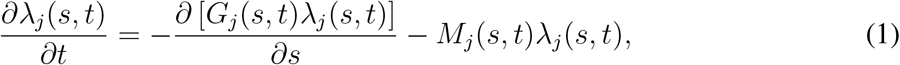

where *λ*_*j*_(*s, t*) is the density of trees of species *j* with size *s* at time *t*, while *G*_*j*_(*·*) and *M*_*j*_(*·*) are species-specific demographic functions describing growth and mortality as a function of size at time *t*, respectively. The first term on the right hand side describes the flux of individuals through the modeled size range, while the second term describes the loss of stems due to mortality. Regeneration is modeled through the boundary condition,

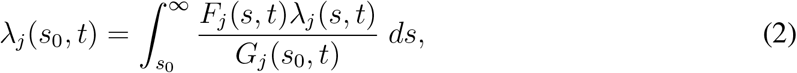

where *s*_0_ is the minimum tree size (in the current context, the smallest diameter stems measured in a forest inventory plot) and *F*_*j*_(*·*) describes specific-specific fecundity as a function of size at time *t*. The term inside the integral describes the species-specific regeneration output from stems of size *s* at time *t* weighted by the initial growth of these stems from the initial size (*s*_0_).

Models of forest dynamics based on the MvF PDE have been used in a range of studies exploring the successional niches of species and plant functional types (Kohyama, 1993; Hurtt et al., 1998; Moorcroft et al., 2001; Strigul et al., 2008; Falster et al., 2011, 2017). In the context of the current study, the MvF PDE provides a computationally efficient, theoretical model of forest dynamics that can be integrated within a broader Bayesian dynamical model. We utilize a hierarchical approach (Fig. 1) to construct our dynamical model defined by data, process, and parameter models (Berliner, 1996).

**Figure 1:**
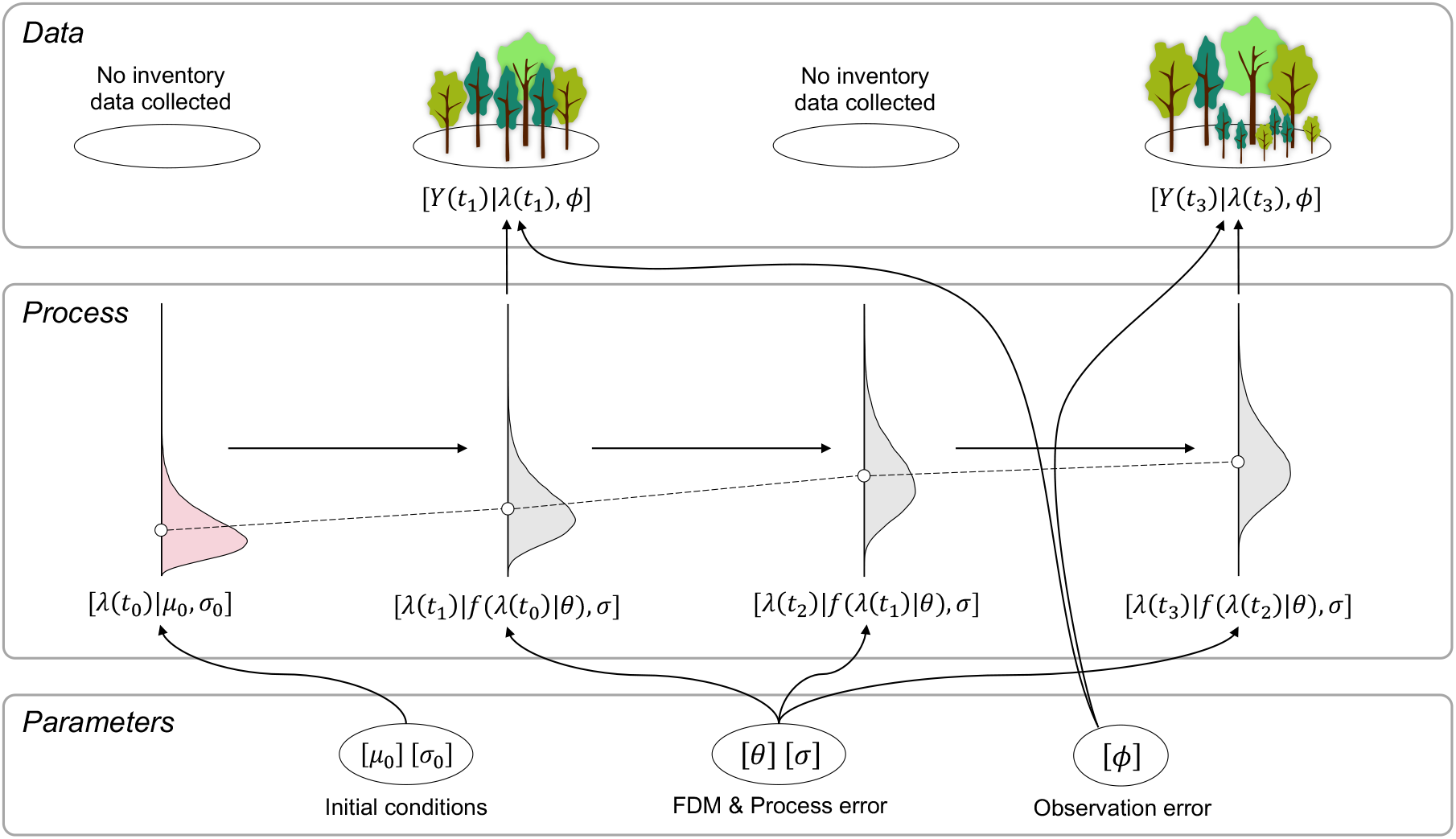
Structure of hierarchical Bayesian dynamical temporal model. Data (*Y* (*t*)) include periodic inventory observations observed within fixed area plots. The process (*λ*(*t*)) represents the latent (unobserved) state of the forest ecosystem driven over time by a model of forest dynamics (*f* ()). Density plots are used to illustrate the fact that a probability distribution exists for each state with points indicating the mean. Parameters include those describing the initial condition of the forest (*µ*_0_, *σ*_0_), model of forest dynamics parameters (*θ*), as well as process (*σ*) and observation (*ϕ*) standard deviations. Square brackets, [], are used to denote probability distribution functions. Arrows indicate dependence among model components. All illustrations were created using Microsoft PowerPoint. Photo credit: Malcolm S. Itter.

#### Data model

We define our data model for plot-level stem counts by species and DBH class consistent with the PEF inventory design prior to 1974. Throughout, we utilize *i* to denote DBH class (*i* = 1, …, *k*), *j* to denote species (*j* = 1, …, *m*), *t* to denote time defined in years (*t* = 1, …, *T*), and 𝓁 to denote inventory plots (𝓁 = 1, …, *L*_*t*_). Note that the number of inventory plots may vary by year. The outcome variable 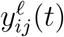 denotes tree counts in DBH class *i* from species *j* observed in plot 𝓁 in year *t*. DBH class counts are modeled using a Poisson distribution,

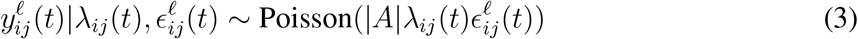

where |*A*| is the size of the plot expressed in hectares, *λ*_*ij*_(*t*) is a rate parameter defining the latent (unobserved) density of trees per hectare in DBH class *i* from species *j* in year *t*, and 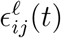 is a multiplicative observation error term accounting for overdispersion driven by plot-to-plot variability within the stand. The latent density *λ*_*ij*_(*t*) represents the discretized size density from the MvF PDE that arises by integrating over stems in the *i*th size class in a given year (*t*),

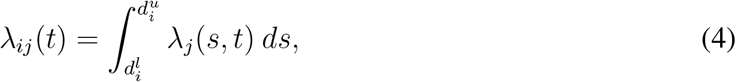

where 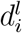 and 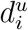 are the lower and upper DBH bounds of the *i*th size class. Both the latent density (*λ*_*ij*_(*t*)) and the observation error term 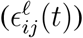 are constrained to be positive. We model the error term using a Gamma distribution,

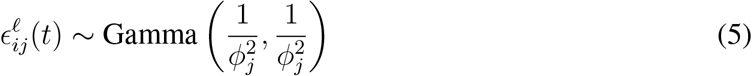

such that 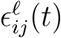 has a mean of 1.0 and variance equal to a species-specific overdispersion parameter 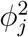. Under the Gamma error model, the marginal distribution for DBH class counts (*y*_*ij*_(*t*)) is negative binomial with mean equal to |*A*| *λ*_*ij*_(*t*) and variance equal to 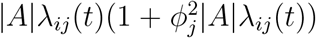 (Lawless, 1987). Hence, the Gamma error model has the computational benefit of allowing us to model DBH class counts conditionally on *λ*_*ij*_(*t*) and 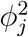 without having to sample the observation error term 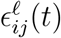 directly. Note, the observation error terms can be obtained through composition sampling following model convergence as described in the Supporting information).

#### Process model

We apply a matrix projection model to predict forest dynamics motivated by the discretization of the MvF PDE in size and time (Eqtn. 4). Matrix projection models refer to a diverse class of population models in which discrete age, size, physiological trait, or life history stage define cohorts used to predict population dynamics in discrete time (Caswell, 2001). When applied in the context of annual forest dynamics with discrete size classes defined by DBH, these models represent a natural extension and formalization of size-class models that have been used for over a century to predict stand-level growth and yield under alternative management scenarios (Weiskittel et al., 2011).

The matrix projection model predicts annual changes in the latent species- and size-specific density (*λ*_*ij*_(*t*)). Consistent with common matrix applications to model forest dynamics (Liang and Picard, 2013), we assume the size-species distribution in year *t* depends only on the distribution in the previous year *t* − 1 (i.e., the temporal evolution of the size-species distribution is a first-order Markov process). The temporal evolution of the rate parameter is modeled as

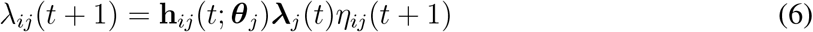

where **h**_*ij*_(*t*; ***θ***_*j*_) is the *i*th row of a (*k* × *k*) species-specific propagator matrix **H**_*j*_(*t*; ***θ***_*j*_) that depends on the year and a set of species-specific demographic parameters ***θ***_*j*_ describing growth, mortality, and recruitment (Supporting information S4). The ***λ***_*j*_(*t*) term denotes a *k*-dimensional vector of latent DBH class densities for the *j*th species: ***λ***_*j*_(*t*) = (*λ*_1*j*_(*t*), *λ*_2*j*_(*t*), …, *λ*_*kj*_(*t*))^⊺^ (where ^⊺^ indicates the transpose). Lastly, *η*_*ij*_(*t* + 1) is a multiplicative process error term reflecting unexplained variation in the temporal evolution of the latent process.

We apply a common parameterization for the propagator matrix (**H**_*j*_) that can be decomposed into separate matrices representing growth, mortality, and fecundity (Liang and Picard, 2013). Specifically, the propagator matrix is given by

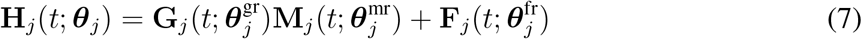

where **G**_*j*_, **M**_*j*_, and **F**_*j*_ are (*k* × *k*) matrices describing growth, mortality, and fecundity across modeled DBH classes for species *j* conditional on demographic rate parameters (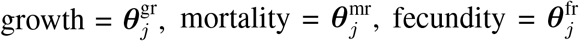, with 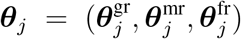. The values of these matrices are estimated using species-specific demographic functions dependent on the defined demographic rate parameters. Here, **G**_*j*_ defines the proportion of individuals in a given DBH class growing into larger DBH classes, **M**_*j*_ defines the proportion of individuals that survive in a given DBH class, and **F**_*j*_ defines the number of individuals that recruit into the smallest DBH class on an annual basis. Consistent with common matrix models for forest dynamics (Liang and Picard, 2013), we assume the Usher property applies such that growth is limited to the next larger size class in a given time step: *i* → *i*+1 for *t* → *t*+1 (note that the 2.5 cm PEF size classes are well above maximum annual growth rates for common Acadian species as modeled in Kuehne et al. (2020)). This assumption combined with the above matrix decomposition leads to a sparse propagator matrix consisting of only (3 × *k*) − 1 demographic rate terms for each species (see Section S3 in the Supporting information).

In the definition of the propagator matrix, we use *t* as a general representation of time-varying conditions that may affect forest demography including exogenous factors such as weather or disturbance and endogenous factors such as density-dependent competition for light, growing space, and other resources within a stand. In the latter case, the propagator matrix depends on the modeled size-species density, ***λ***(*t*), across all species in a stand: ***λ***(*t*) = (***λ***_1_(*t*)^⊺^, …, ***λ***_*m*_(*t*)^⊺^)^⊺^. That is, the propagator matrix accounts for heterogeneous environments affecting demography to the extent that variables describing these environments are included in the applied demographic functions used to define the matrix (see Section S.4 in the Supporting information).

Variability in the temporal evolution of the size-species distribution driven by the propagator matrix is modeled through the process error term (*η*_*ij*_(*t*)). We model the process error term (Eqtn. 6) using a log normal distribution,

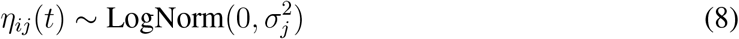

where 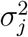 is a species-specific, multiplicative process variance parameter.

#### Demographic functions

In principle, any set of demographic functions may be used to populate the propagator matrix (**H**_*j*_). The only formal constraints are that modeled growth and mortality rates must be between zero and one given that these describe proportions of the population. These functions may come from existing FDMs, knowledge of regional forest systems, or generally applied demographic functions (see Weiskittel et al., 2011, for a thorough review). The parameters of any chosen function are updated based on the inventory data used to inform the Bayesian DSTM. To the extent that functions are taken from previously parameterized FDMs, existing parameter estimates can be used to define prior distributions that help constrain demographic parameters, while allowing for their refinement based on new data, or to fix a subset of parameters focusing statistical information on the parameters to which predictions are most sensitive (Raiho et al., 2020).

We apply a generalized linear modeling approach to estimate demographic rates in the modeled PEF stand based on common link functions. Growth is modeled applying a probit link,

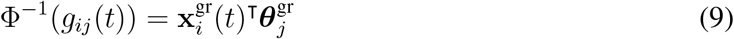

where *g*_*ij*_(*t*) is the size- and species-specific growth rate (the proportion of individuals that grow into the next larger DBH class), Φ is the standard normal cumulative density function, and 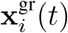 is a vector of variables affecting growth at time *t*. Survivorship is modeled applying a logit link,

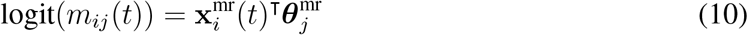

where *m*_*ij*_(*t*) is the size- and species-specific survivorship rate and 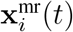 is a vector of variables affecting survivorship at time *t*. We explored a range of predictor variables for growth and survivorship within the modeled PEF stand, but found there was insufficient information to inform parameters beyond simple demographic functions (see Discussion for additional details). We thus estimated growth and survivorship based on single intercept terms, 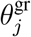 and 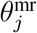 reflecting mean species-specific growth and survivorship rates. The one exception was for eastern white pine, which represents the largest stems in the modeled stand and has greater DBH variability than other modeled species (Fig. S2). We included DBH standardized using a min-max transformation as an additional predictor variable for eastern white pine growth, 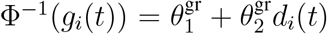, where 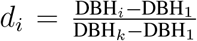, based on exploratory analysis indicating variable growth rates as a function of DBH for the species.

We initially defined a log link function to model fecundity,

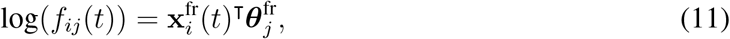

where *f*_*ij*_(*t*) is the size- and species-specific fecundity rate (the per capita number of individuals recruiting into the smallest DBH class) and 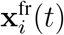 is a vector of variables affecting fecundity at time *t*. However, there were limited observations of regeneration in the modeled PEF stand over the model period. Specifically, ingrowth into the smallest DBH class (2.5 cm) was observed in just three inventory years (1984, 1993, 1999) for two shade-tolerant species (balsam fir, eastern hemlock) in a maximum of two sample plots. Given the limited observations to inform the regeneration component of the process model and the pulsed nature of regeneration events (Díaz-Yáñez et al., 2024), we turned this model component off for the PEF application. Regeneration was instead modeled through the flexible process error term for the smallest size class (*η*_1*j*_(*t*)), which adjusts the latent stem density upward if plot-level observations indicate greater stem counts in the smallest size class than expected based on modeled growth and survivorship rates.

#### Parameter models

The dynamical framework is completed by defining models for demographic rate parameters, initial size-species density values, and observation and process variance parameters. We apply hierarchical priors to model the mean growth and mortality rates among species,

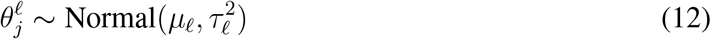

where *µ*_𝓁_ is the mean rate across all *m* species and 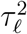 is the among-species variance for the 𝓁th demographic process (𝓁 = 1 for growth, 𝓁 = 2 for mortality). We assign semi-informative normal priors to the demographic rate parameters with means and variances defined to constrain parameters to an ecologically meaningful range based on prior predictive checks under which we simulated data from the dynamical model and compared predictions of overall stem and basal area density to independent, regional stocking guides (Leak et al., 2014). Details on data simulation is provided in the Supporting information. Among species standard deviation parameters (*τ*_𝓁_) are assigned half-Cauchy prior distributions (Gelman, 2006). The hierarchical prior allows for partial pooling of demographic data among species reflecting similarities in the demographic responses of species while allowing for inter-specific variability where it is supported by data. Hierarchical priors can also be helpful when estimating demographic rates of rare or underrepresented species (Dietze et al., 2008).

Initial size-species density values correspond to the latent Poisson rate at the initial time point (***λ***(0)). Modeling initial density values is challenging given the model must balance sufficient flexibility to represent complex size-species distributions, while constraining variability in initial values to reasonable bounds (to prevent numerical instability in the dynamical model). We model initial values using a standardized normal density to represent the size distribution of each species scaled by an estimate of the overall stem density of the species at the initial time point. Specifically,

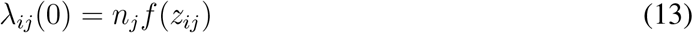

where *n*_*j*_ is the initial stem density of species *j* in terms of trees per hectare and *f* (*z*_*ij*_) is the standardized size density. We apply a standard normal distribution to model the initial size density,

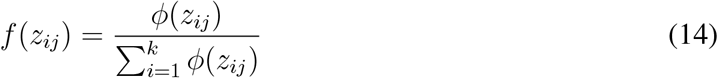

where *ϕ*(*z*_*ij*_) is the standard normal density of *z*_*ij*_, which is defined as 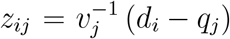 where *d*_*i*_ is the midpoint of the *i*th size class, *q*_*j*_ is the mean of the initial size distribution, and *v*_*j*_ is its standard deviation. Note that the standardized distribution sums to one across the modeled size classes: 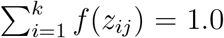 for *j* = 1, …, *m*. Semi-informative log normal and normal priors are applied for the initial density (*n*_*j*_) and mean of the initial size distribution (*q*_*j*_) respectively, with prior mean values defined based on plot-level means of initial inventory observations and prior standard deviations set to 0.05 (on log scale) for the initial stem density and 5.0 cm for the mean of the initial size distribution. Note we use an empirical Bayes approach for the initial density and mean size distribution parameters. We found inventory observations were necessary to help inform initial condition parameters for each species, while prior standard deviations are set to allow for large uncertainty in their values. The initial size distribution standard deviation (*v*_*j*_) is assigned a half-Cauchy distribution. Additional information on prior distributions are provided in the Supporting information.

#### Implementation

We sampled from the joint posterior distribution of the dynamical model parameters using MCMC simulation. Specifically, we applied a Hamiltonian Monte Carlo sampler written in STAN (Stan Development Team, 2024) and implemented using the RStan package (Stan Development Team, 2023) for the R statistical computing environment (R Core Team, 2023). For each version of the model, three MCMC chains were run for a total of 2,000 iterations with a burn-in of 1,000 iterations. Model convergence was assessed based on R-hat values (*<* 1.05), bulk effective sample size (minimum of 100 per chain), and visual inspection of chains for all parameters (Vehtari et al., 2021). Additional information on model implementation including generation of starting values is provided in the Supporting information. Further, we share all model files and associated data processing scripts (Itter and Finley, 2025).

## Results

The modeled PEF stand is even-aged and moves from the stem exclusion to the understory reinitiation stage of development during the 60-year model period (Brissette and Kenefic, 2014). The predicted temporal dynamics of overall stand density reflect these development stages exhibiting characteristic reductions in stem density coincident with increases in basal area (Fig. 2). All species except slow growing, shade tolerant eastern hemlock exhibit decreasing stem densities over the model period contributing to the predicted decreases in overall stem density (Fig. 3). Increases in eastern hemlock density are driven by growth of small diameter stems (2.5-7.6 cm) into the smallest size class depicted (10.2 cm) in Figs. 2 and 3. Fast growing, early successional white pine, which dominates the largest size classes in the stand and eastern hemlock, which dominates intermediate size classes (Fig. 4), account for most of the increase in basal area over the model period (Fig. 2b). The increase in white pine basal area is driven by growth of stems from intermediate size classes (27.9-50.8 cm) into large size classes (53.3-76.2 cm), while the increase in eastern hemlock basal area is predominantly driven by the growth of stems within intermediate size classes (27.9-50.8 cm) over the model period (Figs. 3, 4).

**Figure 2:**
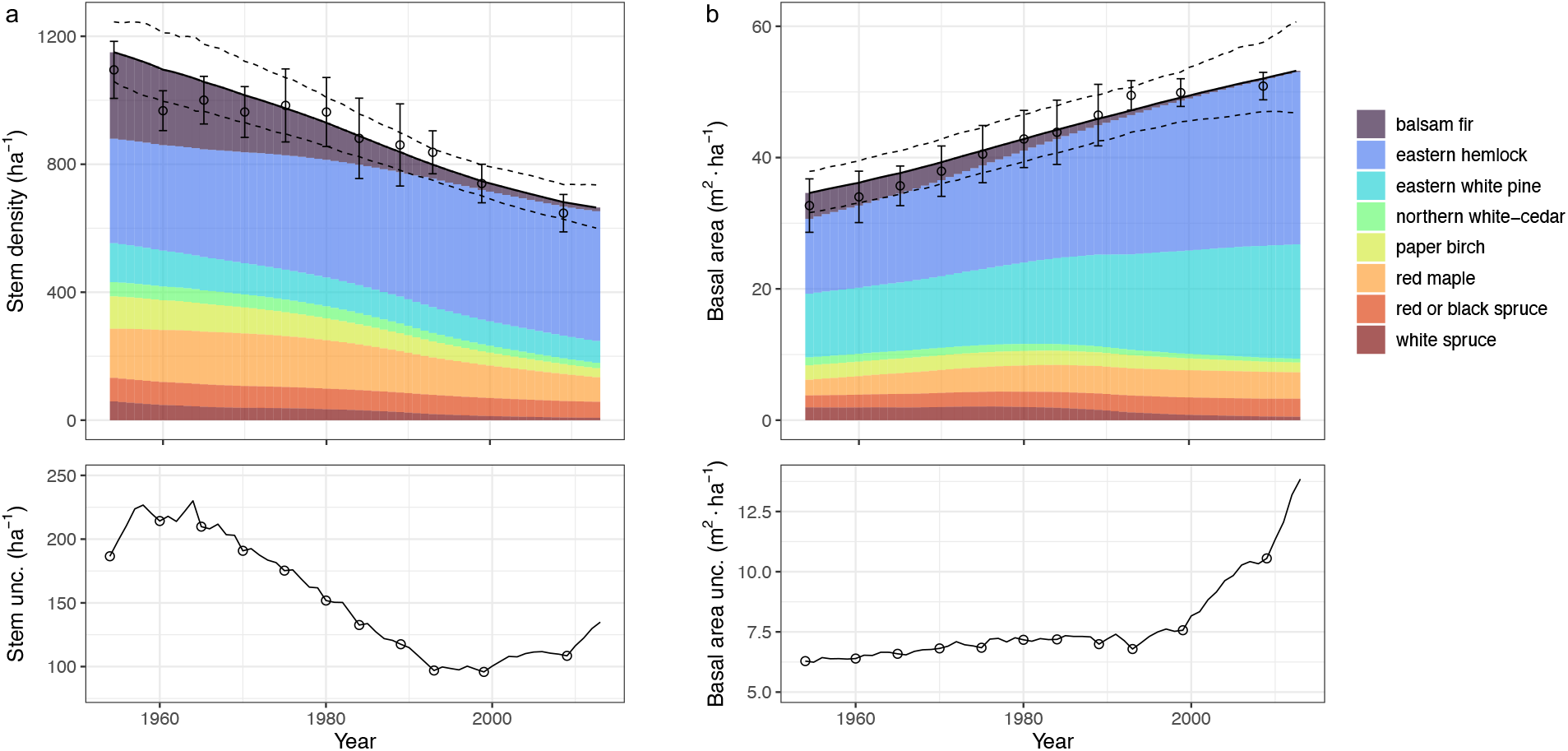
Changes in stand density over the model period (1954-2013) in terms of stems (a), and basal area (b) per hectare. Upper panel represents the posterior mean predicted density values (solid line) and 95 percent credible intervals (dashed lines). Shading indicates the posterior mean predicted density by species. Points represent plot-level observations of stand density with errorbars equivalent to *±*1 standard error. Lower panel represents the 95 percent credible interval width with points indicating inventory years. The lowest size class depicted in 10.2 cm.

**Figure 3:**
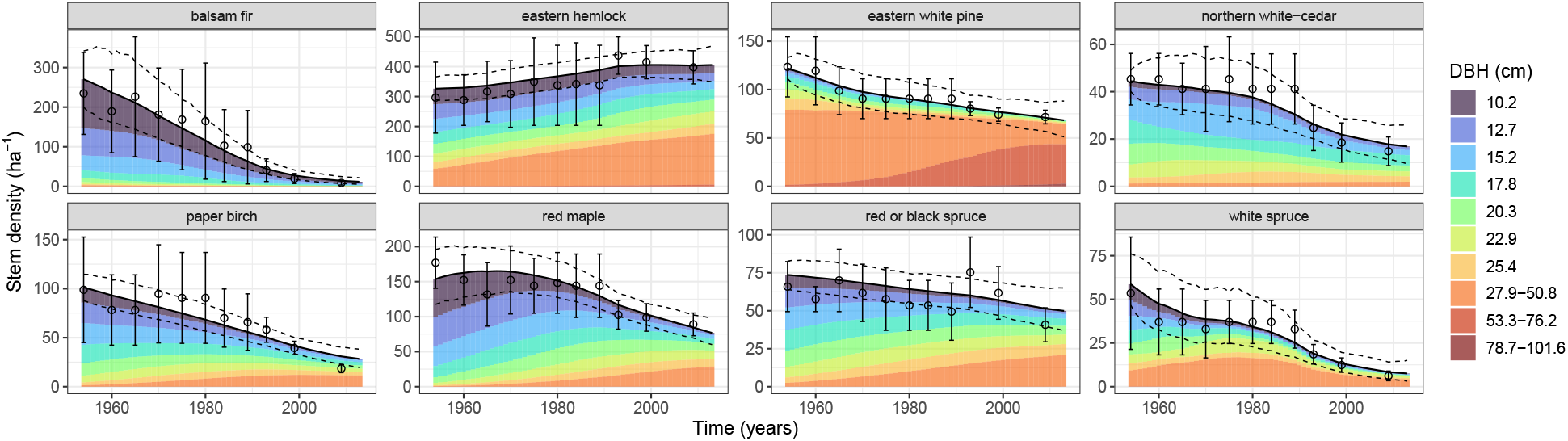
Changes in species-specific stems per hectare over the model period (1954-2013). Solid lines represent the posterior mean predicted stem density (solid line) along with 95 percent credible intervals (dashed lines). Shading indicates the posterior mean predicted density by grouped size class. Points represent plot-level observations of stand density with errorbars equivalent to *±*1 standard error. The lowest size class depicted in 10.2 cm.

**Figure 4:**
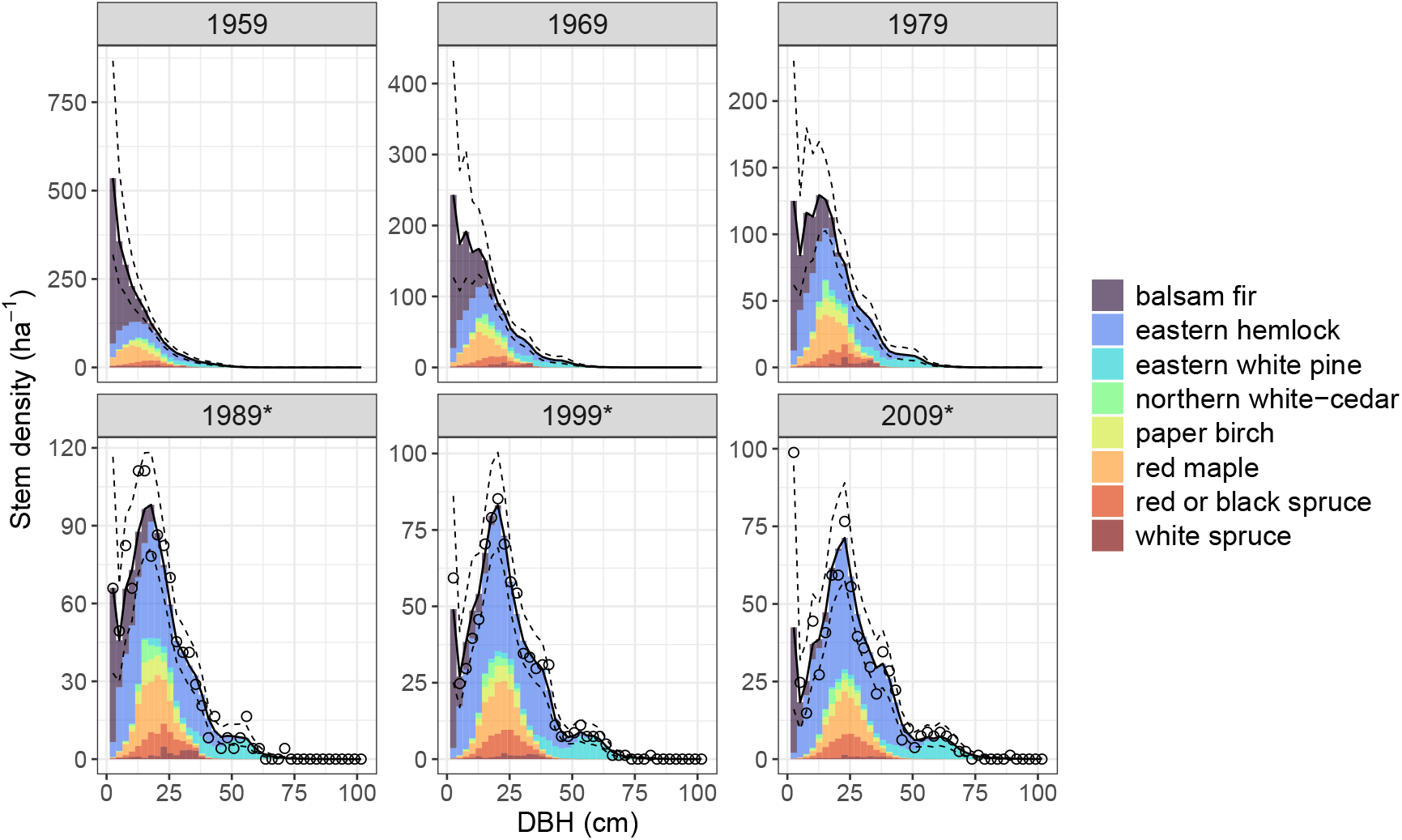
Latent size-species distribution in ten year intervals over the model period beginning in 1959. Solid lines represent the posterior mean predicted total stem density (solid line) along with 95 percent credible intervals (dashed lines). Shading indicates the species-specific posterior mean predicted density within a size class. Points represent plot-level observations of overall stem density by size class.

Other notable species contributing to predicted stand dynamics include balsam fir, red maple, and red/black spruce. Balsam fir is predicted to have high initial stem density, which decreases strongly over the model period (Figs. 2a, 3). The majority of balsam fir stems are predicted to be in the smallest size classes (Fig. 3) with the species dominating these size classes early in the model period (Fig. 4). Both red maple and red/black spruce are predicted to have moderate stem densities and are expected to grow into intermediate size classes over the model period (Figs. 3, 4) contributing to the predicted increase in basal area (Fig. 2b). Changes in the size-species distribution over time reflect a similar pattern to the one described above with the largest size classes composed almost exclusively of white pine with a small proportion of eastern hemlock, the intermediate size classes composed largely of eastern hemlock, and to a lesser extent mid-tolerant red maple and red/black spruce, and the smallest size classes dominated by shade tolerant balsam fir (Fig. 4).

Changes in the composition and size structure of the modeled PEF stand are further reflected in the modeled species-level demographic rates (Fig. 5). Specifically, white pine and red maple have the highest growth rates followed by eastern hemlock, red/black spruce, and paper birch—all species with relatively high abundances in intermediate and large size classes (Fig. 3). Despite its high shade tolerance, balsam fir has the lowest modeled survivorship rate indicative of its predominance in the smallest size classes characterized by high mortality rates. Conversely, shade-tolerant eastern hemlock had the highest modeled survivorship rate, which combined with its moderate growth rate and high stem density helps to explain its large contribution to both the overall stem density and basal area of the modeled PEF stand over time (Fig. 2). All species exhibited relatively low growth rates (less than 12 percent of stems are expected to grow from one size class to the next in a given year) and moderate survivorship rates (at least 90 percent of stems are expected to survive annually; Fig. 5).

**Figure 5:**
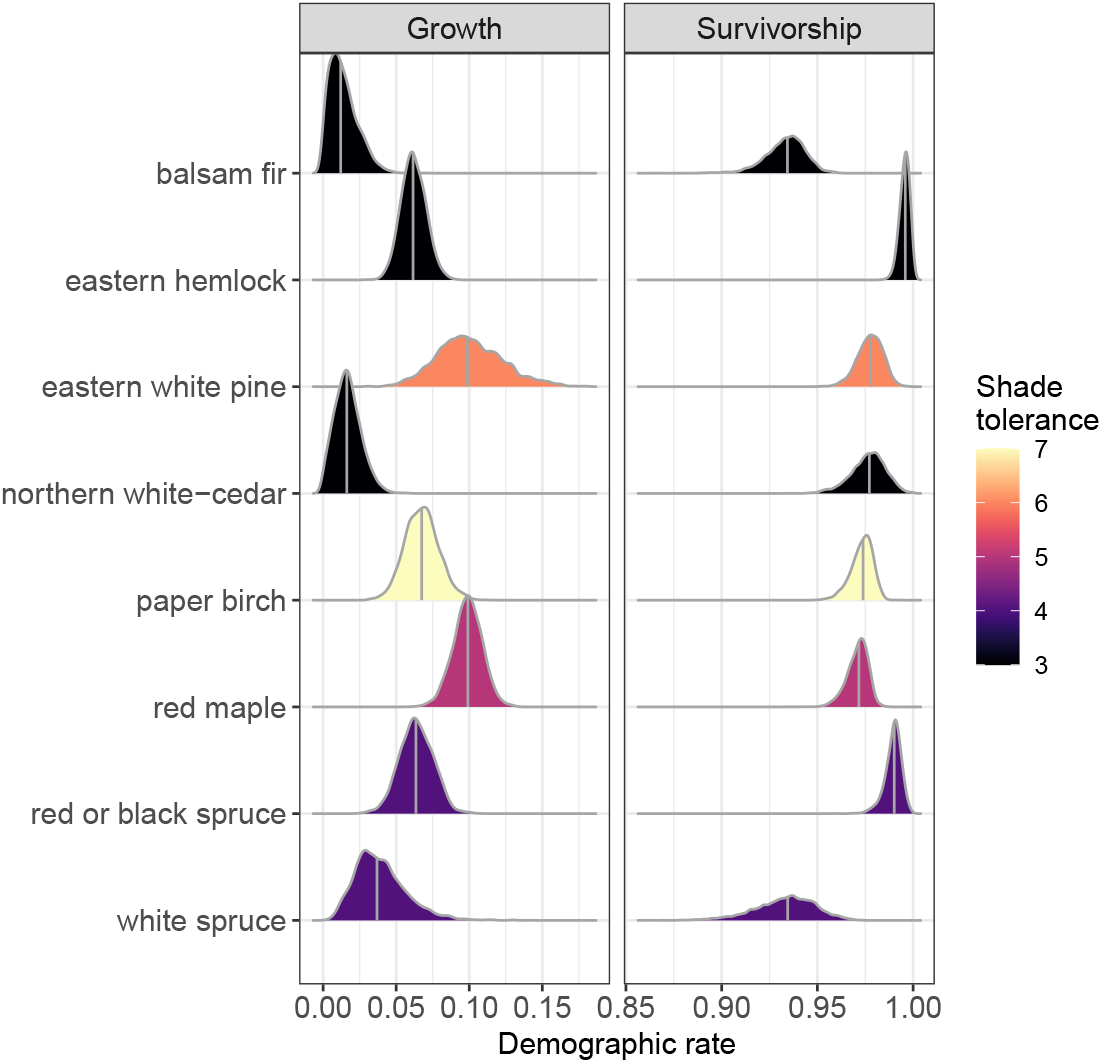
Posterior distributions of modeled species-specific growth and survivorship rates for the Penobscot Experimental Forest. Shading indicates shade tolerance based on index values for northeastern understory plant species with lower values indicating higher shade tolerance (Humbert et al., 2007). Vertical lines indicate posterior median demographic rates. White pine growth rate is shown using a reference diameter at breast height (DBH) value of 25.4 cm.

Uncertainty in predictions of stand dynamics is summarized by 95 percent credible intervals for predictions of overall stem density and basal area (Fig. 2), species-specific stem density (Fig. 3), and the size-species distribution (Fig. 4). These intervals reflect uncertainty in the PEF inventory data (Fig. S3), the process model (Fig. S4), and model parameters (Fig. 5). Figure 2 presents the width of 95 percent credible intervals for predicted overall stem density and basal area with larger values indicative of greater uncertainty. There are clear temporal trends in the uncertainty of both stem and basal area density with the former decreasing and the latter increasing over the model period (Fig. 2). These general trends mirror predicted stand dynamics and are driven by a strong positive association between the posterior mean and standard deviation of latent size-species density values (correlation coefficient equals 0.93), resulting in greater uncertainty for larger predicted density values.

Uncertainty in overall stem density predictions decreases over the first 40 years following a short initial increase, then is stable for approximately 5 years, before exhibiting a slight increase at the end of the model period (Fig. 2a). Uncertainty in overall basal area predictions are more stable with a slight increase over the first 40 years followed by a sharp increase over the last 20 years of the model period (Fig. 2b). While the uncertainty in predicted stem density also increases over the last 20 years, the relative increase is smaller. Interestingly, the increase in uncertainty over the last 20 years of the model period coincides with an increase in the number of permanent sample plots measured per inventory (number of plots increases from three to ten in 1993).

## Discussion

### PEF stand dynamics and associated uncertainty

The goal of the Bayesian dynamical model introduced here is to generate probabilistic predictions of forest states that can be used to make adaptive management decisions accounting for uncertainty in stand-scale forest dynamics. The dynamical model well approximates forest dynamics within the modeled PEF stand. Predicted stand-level changes in composition, size, and density are consistent with expected patterns of Acadian forest development over the stem exclusion and early understory reinitiation phases (Seymour, 2023). Fast growing, early successional white pine forms the main canopy of the stand dominating the largest size classes (Figs. 3, 4). Over time, the initial white pine canopy is predicted to slowly give way to shade-tolerant, slow-growing eastern hemlock, mid-tolerant red maple, and red/black spruce which dominate the midstory of the stand. The understory is dominated by shade tolerant balsam fir and eastern hemlock with the density of stems in the smallest size classes decreasing strongly over time (Fig. 4).

Uncertainty in PEF stand dynamics is expressed through the 95 percent credible intervals of predictions of overall and species-specific stem and basal area density (Figs. 2, 3) as well as predictions of the latent size-species distribution (Fig. 4). The width of these intervals (lower panel in Fig. 2) provides a measure of the posterior variance of predictions reflecting uncertainty in the initial conditions of the modeled stand, the demographic parameters that drive stand dynamics, and the associated process error. The posterior variance further reflects uncertainty in inventory observations, specifically plot-level variability in observed stem densities by species and size class. The effect of inventory observations on predicted stand dynamics can be seen through the reduction in credible interval width in inventory years relative to surrounding years—inventory data constrains the latent stem density (Fig. 2). The amount of reduction in the posterior variance in inventory years is driven by data uncertainty with stronger reductions in the variance when data uncertainty is low (Clark and Bjørnstad, 2004).

The strong association between the posterior mean and standard deviation of latent size-species density values drives changes in the uncertainty of predicted stand dynamics over time. This association relates to the fact that plot-level variability increases as a function of the mean observed stem density (Fig. S3). The applied negative binomial data model helps to account for this relationship given that its variance (data uncertainty) increases as a function of the latent stem density (*λ*_*ij*_(*t*)) and the species-specific overdispersion parameter 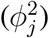 allowing for additional plot-level variability. Increases in plot-level variability (data uncertainty) for larger observed stem densities helps to explain the trends in stem density and basal area uncertainty (Fig. 2). The beginning of the model period coincides with the stem exclusion phase of development characterized by high stem densities in small size classes. The large plot-level variability associated with these high observed stem densities (high data uncertainty) contributes to a poorly constrained latent stem density (Figs. 2a, S6). The two species with the largest stem densities in this period, balsam fir and eastern hemlock (Figs. 3, 4), have some of the largest overdispersion parameters among all species indicating additional plot-level variability in observations (Fig. S3). As observed stem densities decrease over time (driven by density-dependent mortality), so does the associated data uncertainty, which contributes to reduced uncertainty in the latent stem density. Balsam fir plays an outsize role in these dynamics given it has the highest overdispersion parameter of any species (Fig. S3) and high initial stem densities, which decrease to almost zero over the model period (Fig. 4).

Trends in basal area uncertainty are similarly driven by uncertainty in the latent stem density, except that basal area uncertainty is more sensitive to variability in larger size classes given their greater relative contribution to the overall basal area. This helps to explain the stability in the uncertainty of predicted basal area over the first 40 years of the model period when the largest predicted stem densities are in small size classes, which contribute little to the overall basal area and its associated uncertainty (Fig. 2b). As the predicted stem density in intermediate to large size classes increases later in the model period, driven by the growth of eastern white pine and eastern hemlock (Figs. 3, 4), so does the uncertainty in predicted basal area (Fig. 2b).

The positive correlation between mean observed stem density and associated plot-level variability further helps to explain the counter-intuitive increase in the uncertainty of predicted stand dynamics over the last 20 years of the model period when the number of permanent sample plots increases from three to ten. While we would expect the uncertainty in stand dynamics to decrease as a function of sampling intensity, we observe the opposite within the modeled PEF stand. In this case, the increase in uncertainty relates to stem density increases in intermediate and large size classes dominated by eastern hemlock and white pine, respectively, which have a large effect on total basal area and its associated uncertainty (Figs. 3, 4). In particular, there is a large increase in the mean observed stem density and associated plot-level variability of eastern hemlock in the 27.9 to 50.8 cm DBH classes with the addition of new permanent sample plots in 1993 (Fig. S6). At this time, eastern hemlock is the dominant species in terms of both stem density and basal area (Fig. 2) with the majority of stems in the noted DBH class range (Fig. 3) making the model particularly sensitive to variability in this species-size combination. In addition, the ten year gap between forest inventories in 1999 and 2009 due to a missed five-year re-measurement in 2004 (Table S1) further contributed to the increased uncertainty over the last 20 years of the model period.

### Inventory data uncertainty

As highlighted above, plot-level variability in observed stem densities by species and size class (data uncertainty) contributes strongly to the uncertainty in predicted PEF stand dynamics. This variability reflects heterogeneous stand conditions, particularly in the density of stems in the smallest size classes dominated by balsam fir and eastern hemlock (Fig. S6). Increasing the number of permanent sample plots included in inventories generally helps to reduce the variability in estimates of mean stand conditions (Burkhart et al., 2018). This is true for estimates of the overall PEF stand density. Note that the standard errors of mean observed stem density and basal area are lower in the last three inventory years when there are ten instead of three permanent sample plots (Figs. 2, 3). Despite this, there is an increase in the uncertainty of predicted stand density over this period. This is due to the fact that the dynamical model predicts changes in the size-species distribution rather than the overall stand density and, as such, is sensitive to large variability in individual species-by-size classes (in this case eastern hemlock in 27.9 to 50.8 cm DBH classes; Fig. S6). An alternative sampling strategy to reduce variability in stand density estimates is to increase the size of plots (Burkhart et al., 2018). This may be a more effective strategy to reduce data uncertainty in the modeled PEF stand given that the smallest size classes tend to have the greatest variability (Fig. S6) and are measured using the smallest plot sizes (0.008 or 0.02 ha).

Beyond modifying the structure and/or number of inventory plots, the impact of heterogeneous stand conditions on the uncertainty of predicted stand density could be reduced by modeling forest dynamics at the plot level (i.e., decreasing the spatial scale at which forest dynamics are modeled). The choice to model forest dynamics at the stand-scale relates to our goal of ultimately applying the dynamical model to help inform adaptive management decisions. The stand is the inferential scale of interest given its connection to management decisions and forest resilience to global change (Seidl et al., 2013). In this context, data uncertainty related to heterogeneous stand conditions is not necessarily bad. Rather, it provides an accurate and objective measure of the variability in the underlying stand conditions and the potential response of the system to future management, consistent with the goals of any forest inventory (Burkhart et al., 2018). Note that while the process model operates at a stand scale, the dynamical model still allows for plot-level predictions of stem density and basal area through the data model (Figs. S7, S8).

### Improving predictability of forest dynamics through model refinement

The dynamical model applies simple demographic functions and a simple initial condition model to describe changes in the latent size-species distribution over time. Despite its simplicity, process error standard deviations are relatively low for all species (Fig. S4) leading to small process error values (mean process error values range from 0.9 to 1.3) suggesting that the model approximates observed PEF stand dynamics reasonably well (a process error value of 1.0 indicates no error). The predictive performance of the simple demographic functions is likely due to the slow rates of change within the modeled PEF stand. Note that the observed size range of each species is nearly constant over the model period with the exception of eastern white pine and to a lessor extent eastern hemlock and red maple (Fig. S2) leading to low modeled growth rates (Fig. 5). The biggest change in the modeled PEF stand is a consistent decline in stem density (Fig. 3) driven by density-dependent mortality occurring during the stem exclusion stage of development, which contributes to moderate annual survivorship rates (Fig. 5).

We do not expect the applied demographic functions and initial condition model to perform well generally. They are too simple and do not integrate sufficient ecological detail to make accurate predictions for a broad diversity of sites and conditions. This is an important limitation given that a goal of the model is to predict forest dynamics under global change. In general, process models that closely approximate the underlying mechanisms of ecosystem dynamics are thought to provide better predictions under novel future conditions (Yates et al., 2018). The specific demographic functions and initial condition model applied in the PEF analysis should not be viewed as representative of the mechanisms underlying forest dynamics. Rather, they reflect a compromise between model complexity (ecological detail) and parameter identifiability (Luo et al., 2009). We initially defined more complex demographic functions and a more flexible initial condition model that we expect would better approximate general forest dynamics (see Supporting Information). For example, initial demographic functions modeled growth as a concave function of size modified by density-dependent competition and mortality as a function of the modeled growth rate. Unfortunately, there were significant identifiability issues when applying these complex functions, leading to the simple functions applied here.

Several factors likely contributed to the lack of identifiability in the more complex demographic functions and initial condition model. Without independent information, we found it necessary to connect the initial condition model to the first set of inventory observations (i.e., initialize the model in the first year inventory data was collected). With only three permanent sample plots measured in the first inventory, there was limited information to inform the initial condition parameters. The large variability among these initial plots given high stem counts in small size classes further exacerbated this issue. Higher sampling intensity in the initial inventory year may have allowed for a more flexible model that better approximates the initial size-species distribution. Note that while it is common to apply a model “spin-up” approach to simulate initial conditions when modeling forest dynamics (Raiho et al., 2020), we did not have a parameterized FDM to predict initial conditions in the absence of inventory data.

The identifiability of demographic parameters was affected by the limited observed size range for model species (Fig. S2). Most species tended to be present in only a small subset of size classes over the model period experiencing a similar level of size-structured competition making it challenging to parameterize species-specific, non-linear demographic functions including size and competition terms. Further, there were almost no observations of ingrowth during the model period such that no information was available to estimate fecundity parameters. Identifiability was also affected by strong correlation among modeled demographic parameters. Correlation among demographic parameters has been found in previous studies (Wilson et al., 2019) and is not surprising given the expected dependence among forest demographic processes (Cailleret et al., 2020). A benefit of the Bayesian dynamical model is that it estimates these demographic parameters jointly, allowing for their correlation to be explicitly modeled. However, estimating this correlation structure and identifying correlated parameters can be challenging for MCMC routines and usually requires a large amount of data (Roberts and Sahu, 1997). Importantly, strong correlations may also be due to redundant or confounded parameters, which demand that demographic functions be simplified or reparameterized (Betancourt, 2018). Existing approaches to parameterize FDMs including the use of restricted maximum likelihood estimation applied separately for each species and demographic process do not account for these correlations, potentially leading to overspecified models that provide biased predictions.

Refinements to the dynamical model aimed at better representing the processes underlying forest dynamics include applying demographic functions that integrate measures of density-dependent competition and climate. Representing competition at the stand (or population) scale is challenging, but critical to the ability of the dynamical model to accurately represent stand dynamics. While simple measures such as “basal area larger” have historically been used in stand-level models (Weiskittel et al., 2011), alternatives such as the perfect plasticity approximation have been shown to better represent resource competition (Strigul et al., 2008). Existing measures of stand-level competition all assume homogeneous competitive conditions throughout the stand. Given the importance of spatial heterogeneity to forest resilience and adaptive management (Wikle and D’Amato, 2023), future work should focus on developing approximations of the mean competitive growing environment that better account for variable stand conditions. Demographic functions should also incorporate climate as a driver of forest dynamics, particularly given interest in applying the dynamical model to predict forest responses to global change. Climate-derived variables (e.g., vapor pressure deficit) can be incorporated as modifiers to baseline demographic rates similar to existing FDMs (Bugmann, 1996), or they can be used to predict local demographic parameter values along with physical site characteristics as part of a broader spatio-temporal model (Finley et al., 2009). Synthesizing tree-ring data with long-term forest inventory observations may allow for inference on the effects of climatic variables on forest demographic rates (Itter et al., 2017; Heilman et al., 2022).

Additional model refinements include the integration of more complex process and observation error models. While the initial dynamical model assumes an independent error structure for both the process and observation error (Eqtns. 5, 8), there may be additional dependence in these error terms, which should be accounted for through modeled covariance among species and size classes. Given the relatively high dimension of the size-species distribution, it is likely that some form of dimension reduction will be required to model these covariances. The process error variance may also vary over time (Raiho et al., 2020). While flexible modeling approaches exist for dealing with high-dimensional, time varying error structures (Taylor-Rodriguez et al., 2019), they require a large amount of data to fit. As such, we did not attempt to model these complex error structures in the initial dynamical model.

### Model extensions

The lack of inventory data reflecting a range of growing conditions including well-replicated observations of each species’ growth, mortality, and fecundity under variable environments and stand conditions is the biggest obstacle to defining a richer dynamical model for the PEF (as described above). Future work will focus on extending the dynamical temporal model defined here to a dynamical spatio-temporal model informed by regional forest inventory data. While previous studies have modeled spatio-temporal forest dynamics using structured population models based on the MvF PDE (Kohyama, 1993; Falster et al., 2017), none have directly linked these population models to data within a Bayesian DSTM that supports uncertainty quantification and propagation. Such a model would provide probabilistic predictions of stand-scale forest dynamics over broad spatial extents.

Additional work will focus on integrating natural disturbance and forest management as a driver of forest dynamics within the dynamical model. The integration of natural disturbance is essential to predict forest dynamics under global change. Incorporating management as a driver of forest dynamics will allow for probabilistic predictions of forest responses to adaptive management strategies to assess their capacity to promote forest resilience and adaptation to novel climate and disturbance regimes. These predictions can be integrated within a Bayesian decision theoretic framework to identify optimal management strategies that formally account for the uncertainty in predicted forest responses (Dorazio and Johnson, 2003). While an initial framework for integrating forest management into the dynamical model has been defined (Itter et al., 2025), additional work remains to scale this framework across space and allow for interactions with climate and natural disturbance.

## Conclusion

The Bayesian dynamical model presented here provides an initial probabilistic framework for predicting stand-scale forest dynamics informed by inventory data. The MvF PDE serves as the theoretical foundation of the model framework providing process-based representation of size-structured population dynamics based on relatively few parameters. While the demographic functions applied are simple, the dynamical framework is extremely flexible allowing for varying levels of ecological detail to be integrated into the model so long as there is sufficient information in applied data to support it. In this context, we view the identifiability issues encountered in the PEF application as a strength of the modeling approach limiting potential overparameterization. The initial dynamical model represents a first step towards a spatio-temporal model providing probabilistic predictions of stand-scale forest dynamics across broad spatial extents under novel climate and disturbance regimes. As such, it represents an important innovation in the pursuit of applied understanding of forest responses to global change and informed adaptive management.

## Acknowledgments

This study was supported by funding from the National Science Foundation grant numbers DEB-1946007 (MSI, AOF) and DEB-2213565 (AOF). Work was further supported by the USDA Forest Service grant number 23-JV-11242305-080 (MSI) and its partnership with the National Council for Air and Stream Improvement, Inc. (AOF).

