## Supporting information for "Making more with forest inventory data: Toward a scalable, dynamical model of forest change"

**Title:** Toward improved uncertainty quantification in predictions of forest dynamics: A dynamical model of forest change

### S1 Supplemental tables and figures

| Inventory years | Number of plots |
|---|---|
| 1954 | 3 |
| 1960 | 3 |
| 1965 | 3 |
| 1970 | 3 |
| 1975 | 3 |
| 1980 | 3 |
| 1984 | 3 |
| 1989 | 3 |
| 1993 | 10 |
| 1999 | 10 |
| 2009 | 10 |

Table S1: Number of permanent sample plots in each inventory year within management unit 32B of the Penobscot Experimental Forest.

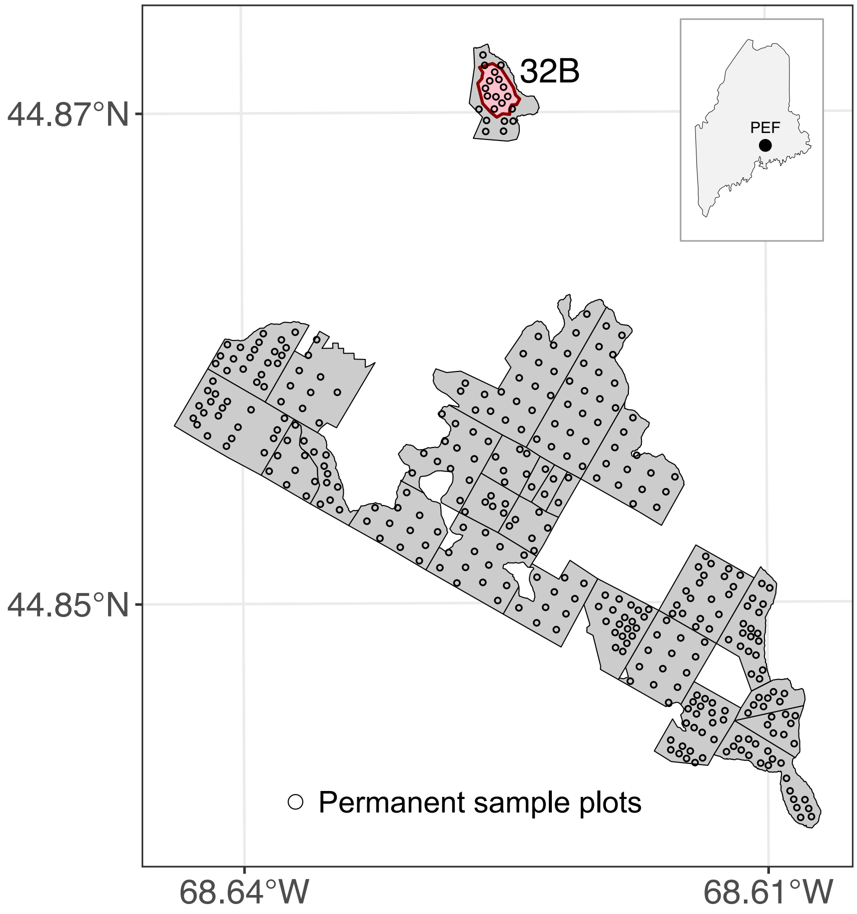


Figure S1: Map of the Penobscot Experimental Forest management units with the modeled unit (32B) highlighted. Points indicate permanent sample plot locations.

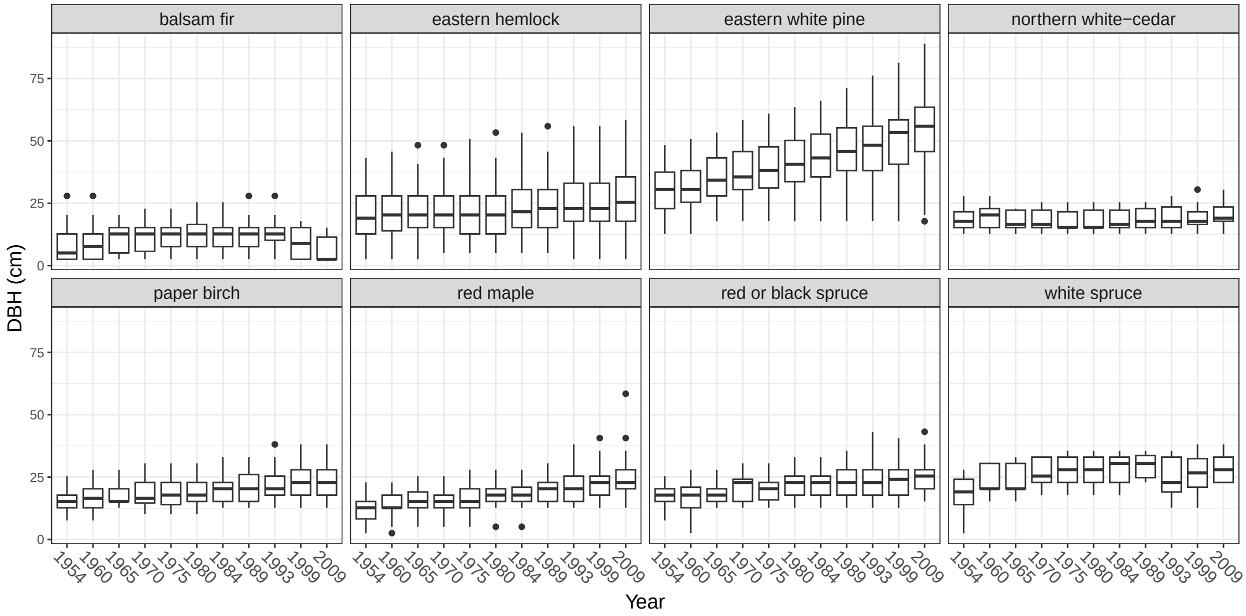


Figure S2: Boxplots of the observed tree diameter classes by species in each inventory year pooled across measured plots (sample size varies by year as indicated in Table S1).

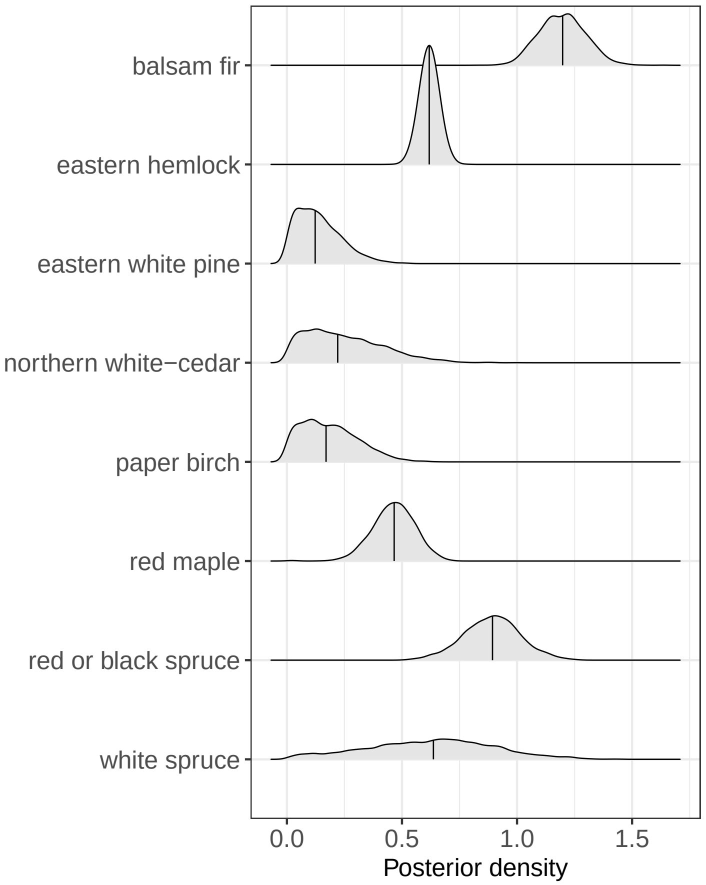


Figure S3: Posterior distribution of the species-specific observation error standard deviation equivalent to the square-root of the overdispersion parameter ($\phi_j$).

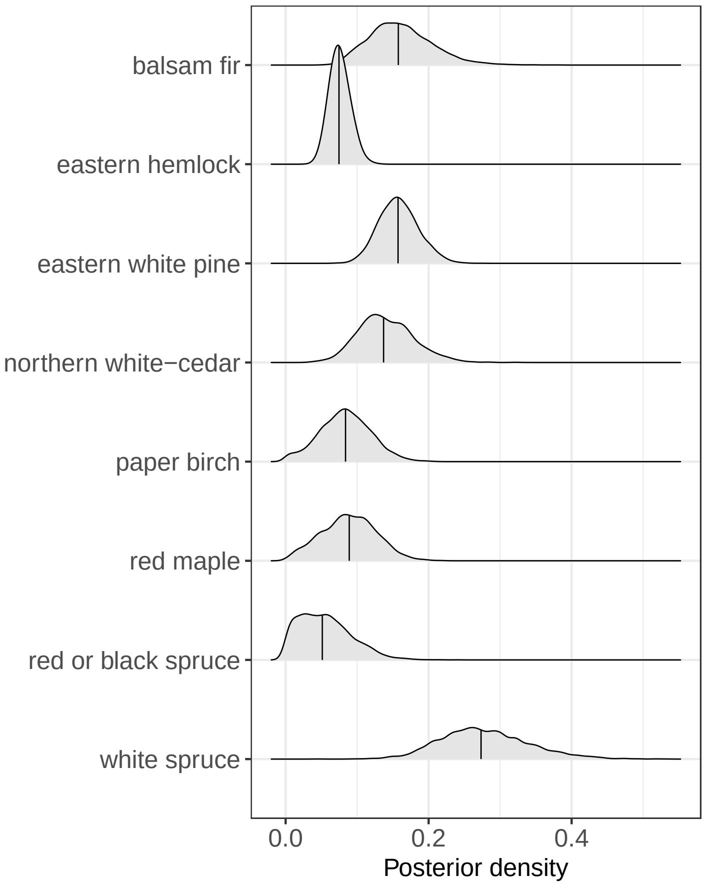


Figure S4: Posterior distribution of the species-specific process error standard deviation ($\sigma_j$).

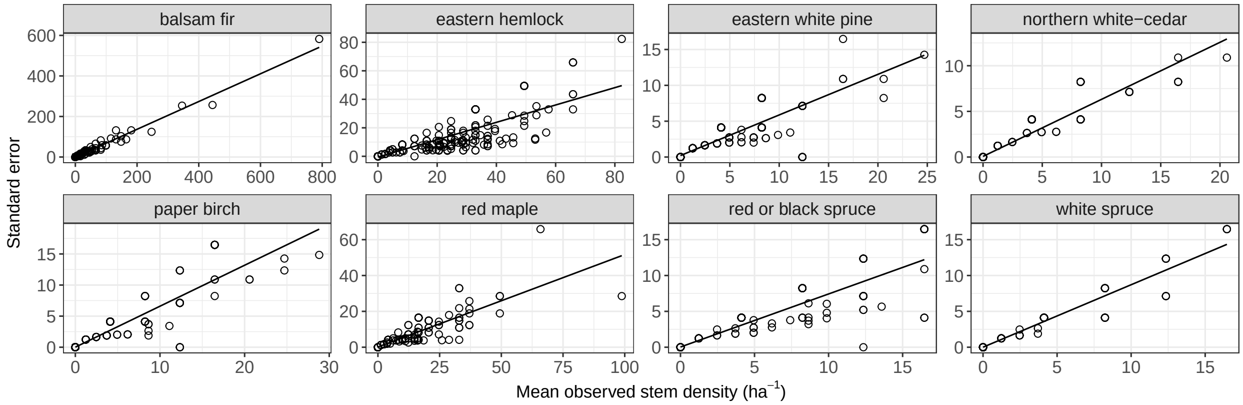


Figure S5: Plot-level standard error versus mean observed stem density by species. Points correspond to individual size classes in each inventory.

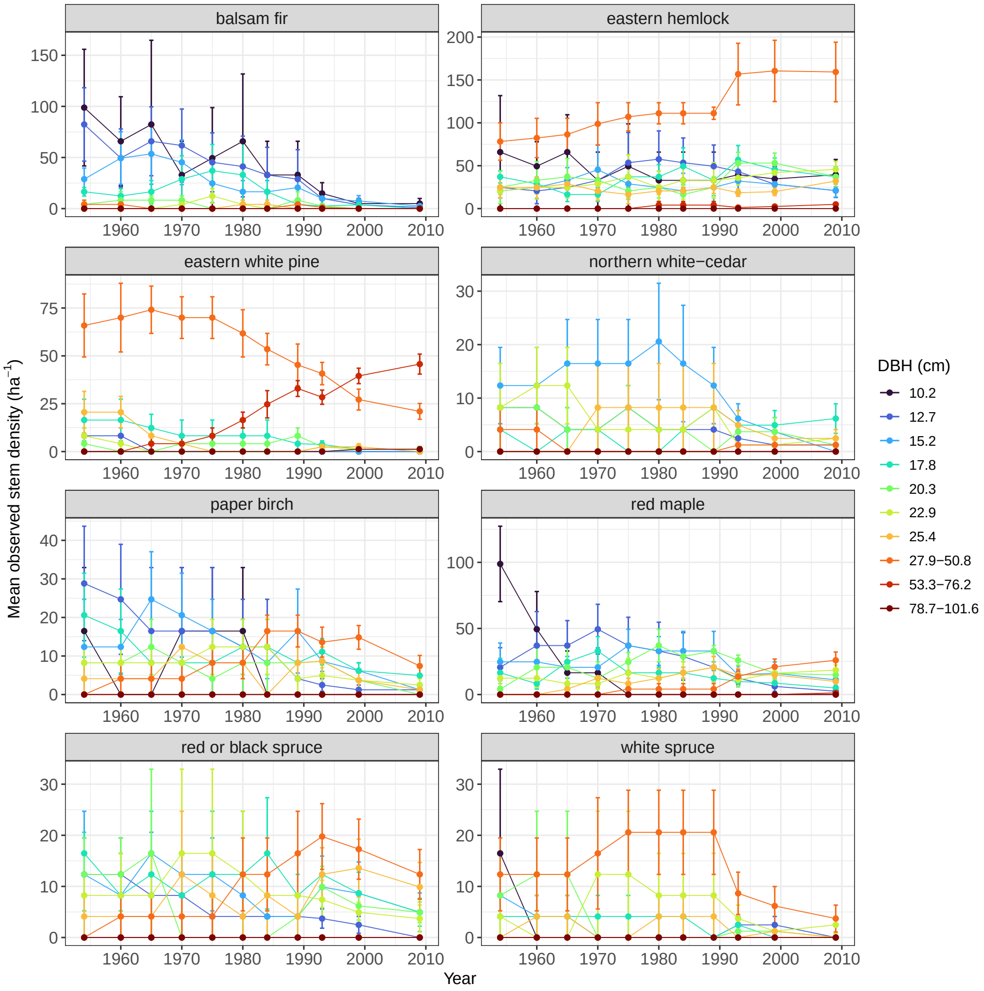


Figure S6: Mean observed stem density in grouped size classes over time by species. Error bars indicated ± 1 standard error.

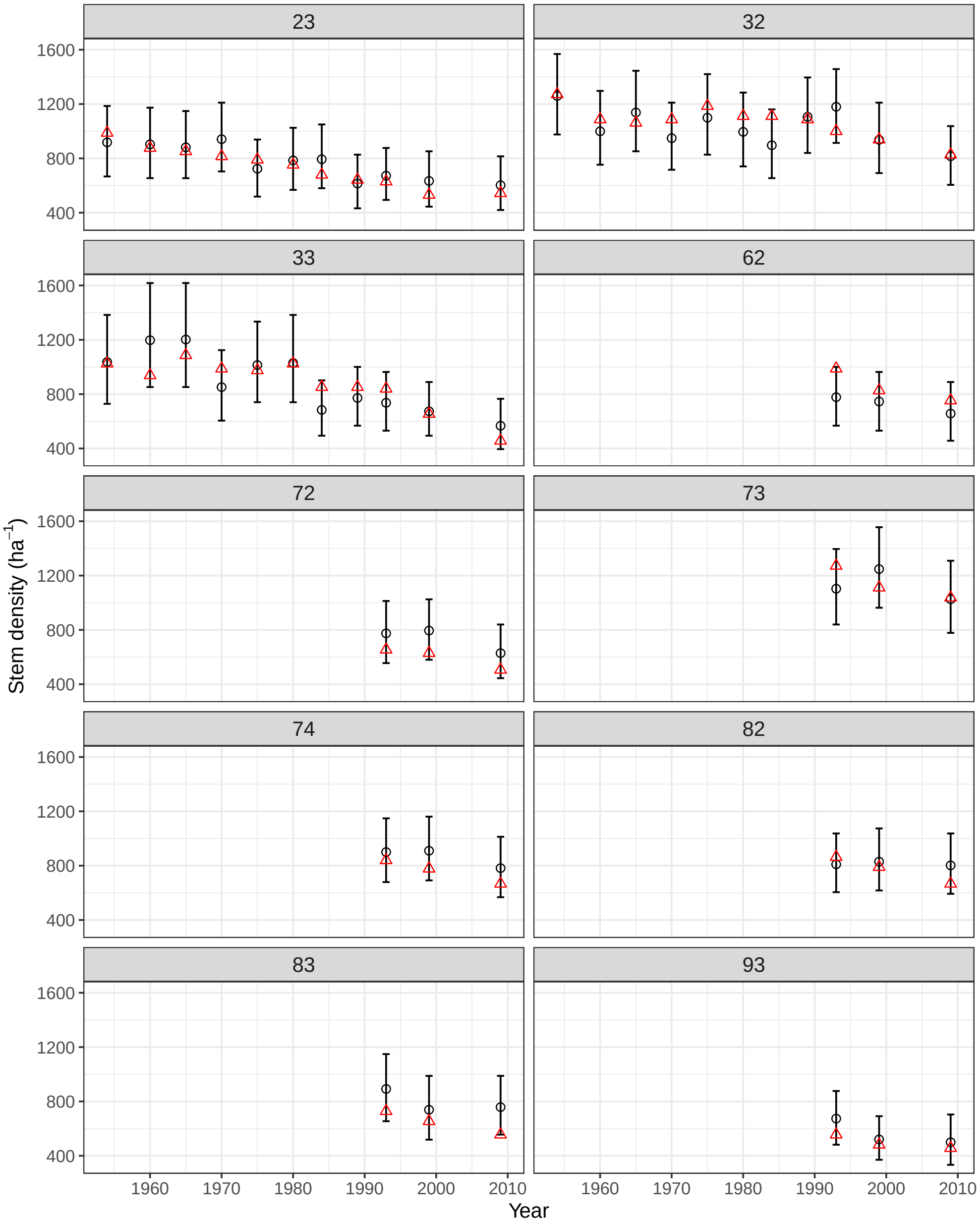


Figure S7: Plot-level predictions of stem density for each inventory year relative to observed values. Open circles with error bar correspond to posterior mean predicted density and associated 95 percent credible interval bounds. Red triangles indicate the observed density. Note that seven inventory plots were measured only in the last three inventories.

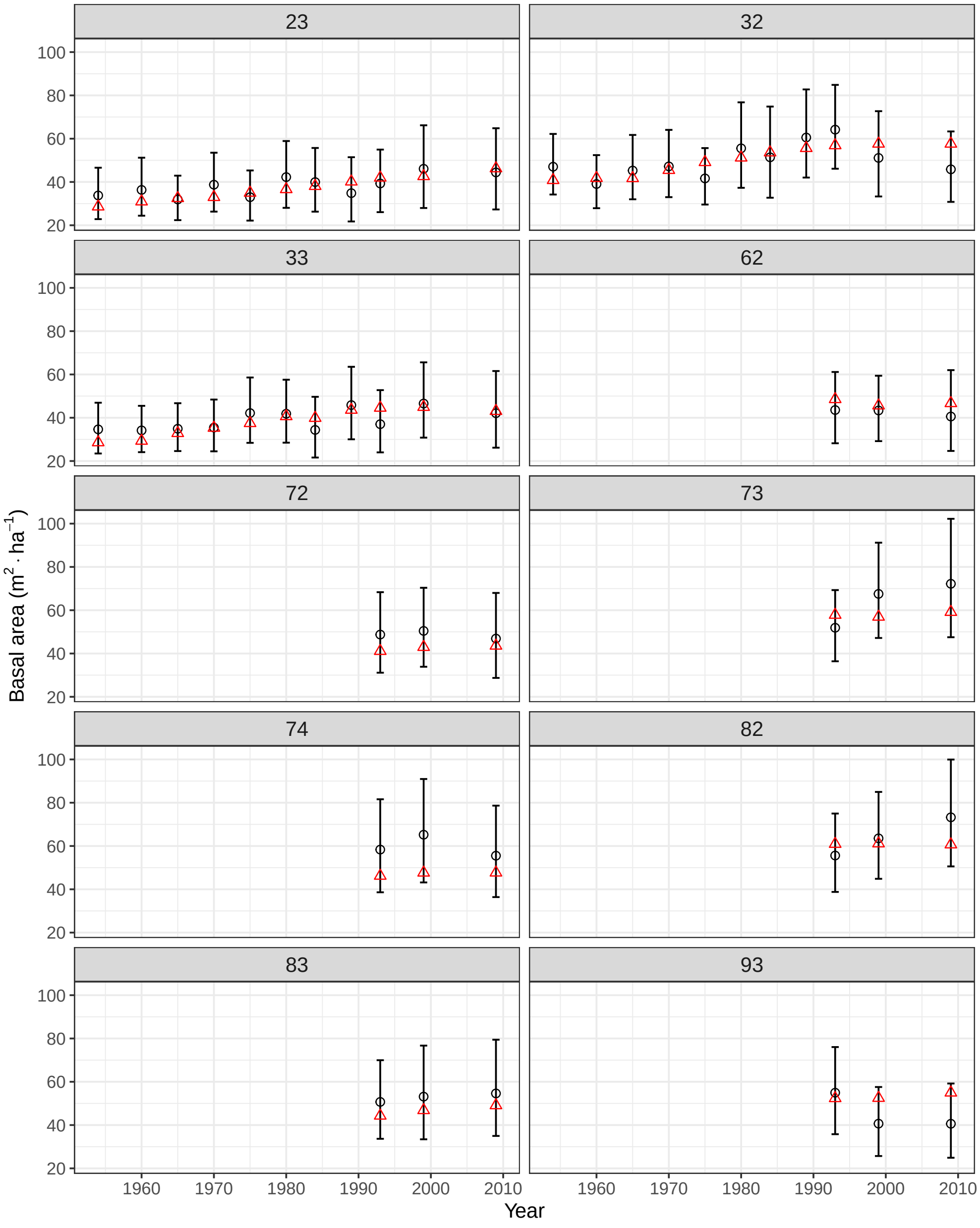


Figure S8: Plot-level predictions of basal area density for each inventory year relative to observed values. Open circles with error bar correspond to posterior mean predicted density and associated 95 percent credible interval bounds. Red triangles indicate the observed density. Note that seven inventory plots were measured only in the last three inventories.

#### S2 Simulated eastern temperate forest succession

We simulated eastern temperate forest succession over 100 years based on the dynamical temporal model defined in the main text. Successional dynamics were simulated for a three species system representative of different functional groups including a fast growing, shade intolerant "pioneer" species, a mid-tolerant, slower growing "gap-phase" species, and a slow growing, "shade-tolerant" species. The demographic functions used to simulate long-term forest dynamics are defined below (Section S4.2). Demographic parameters were set to reflect the different functional groups. Initial conditions were set to approximate a forest stand nearing the end of the establishment phase of development. We further included a single process error standard deviation for all three species. The long-term dynamics of the simulated eastern temperate forest are illustrated by the evolution of the size-species distribution over time (Fig. S9) as well as changes in the species-specific stem and basal area density over time (Fig. S10). The modeled dynamics indicate an initial main canopy composed of the pioneer species with the gap-phase species located in an intermediate canopy position and the shade-tolerant species dominating the understory. Over time, the pioneer species is slowly replaced in the canopy by the gap-phase species while the shade-tolerant grows into the midstory. At the end of the model period, the main canopy is composed of a mixture of primarily pioneer and gap-phase individuals with shade tolerant individuals dominating the midstory and understory.

We fit the dynamical model to simulated CFI observations for ten 0.08 ha plots remeasured every five years over the 100-year simulation period according to the data model (Eqtn. 1), in order to check the model's ability to recover underlying demographic and uncertainty parameters. We fixed parameters associated with the basal area larger competition term due to limited variability over the model period. Further, we fixed mortality parameters associated with a quadratic size term (included to approximate increasing mortality as trees reach their maximum size) given limited densities of individuals in maximum size classes. All model parameters were well identified with 95 percent credible intervals capturing the simulated parameter value.

The R code to generate the simulated data and fit the dynamical model is included as part of the supporting information. Interested users can modify the simulation parameters as well as the demographic functions and parameters used to generate the data to explore alternative temporal patterns of forest change.

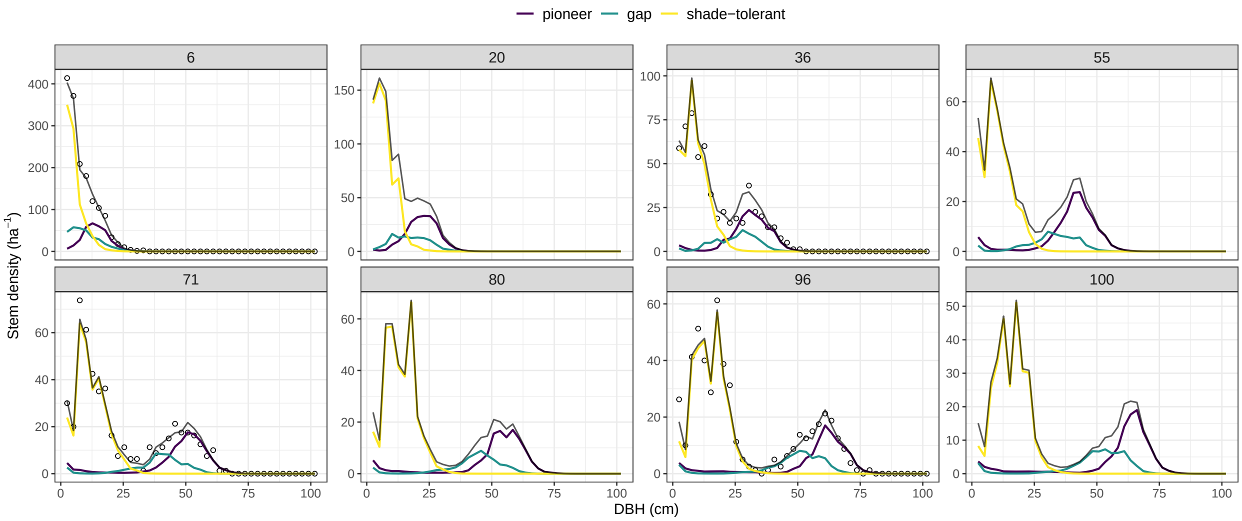


Figure S9: Predictions of the simulated size-species distribution in eight years during the 100-year simulated model period. Grey line is the latent overall size density. Points are the mean overall size density across simulated CFI plots (only simulated for inventory years). Colored lines are the size density by species.

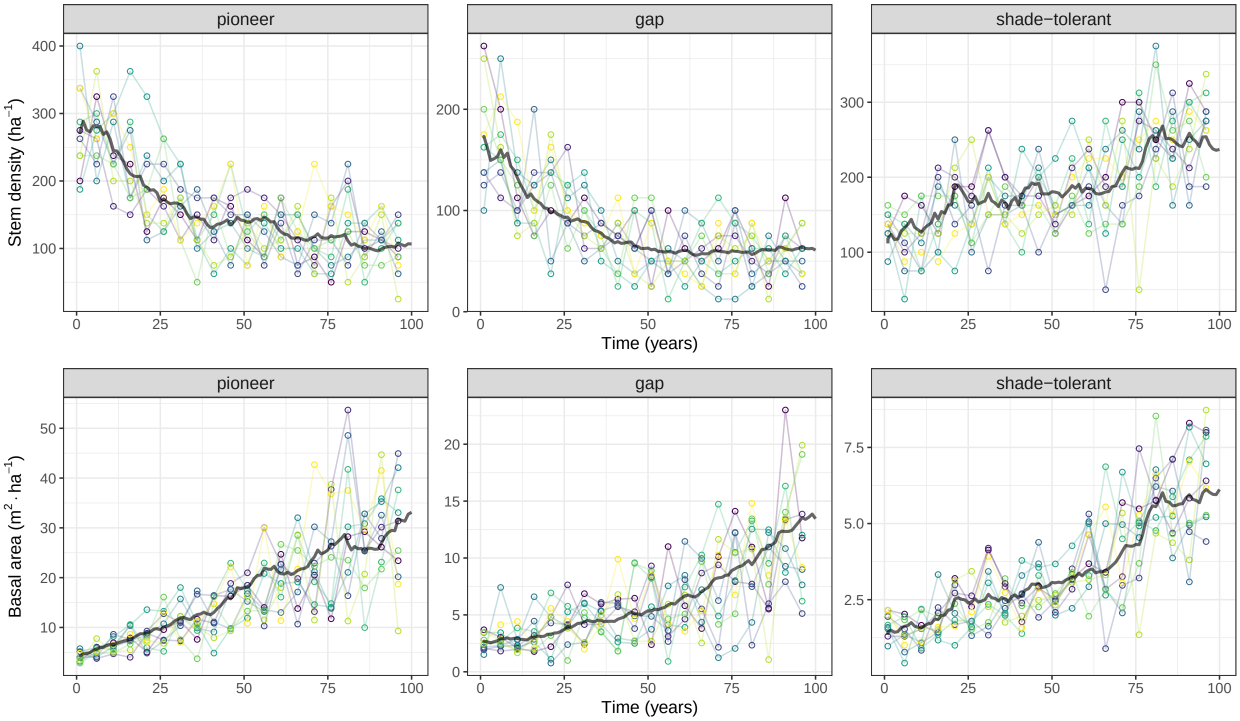


Figure S10: Changes in the simulated species-specific density in terms of stems and basal area per hectare over the simulated model period. Grey line is the latent density. Points are periodic (every five years) observations from ten simulated CFI plots denoted by different colors and connected for reference.

#### S3 Matrix projection model

As noted in the main text, the matrix projection model motivated by the discretization of the McKendrick-von Foerster partial differential equation (MvF PDE) results in a sparse, ecologically-meaningful representation of the complex dynamics of the size-species distribution within a forest stand over time. Expanding the process model (Eqtn. 4) for each size class of a given species $j$ (temporarily dropping the process error term for simplicity) results in the following set of equations,

$$\mathbf{h}_{ij}(t;\boldsymbol{\theta}_j)\boldsymbol{\lambda}_j(t) = \begin{cases} m_{ij}(t)(1-g_{ij}(t))\lambda_{ij}(t) + \sum_{\ell=1}^{k} f_{\ell j}(t)\lambda_{\ell j}(t) & i=1 \\ m_{ij}(t)[g_{(i-1)j}(t)\lambda_{(i-1)j}(t) + (1-g_{ij}(t))\lambda_{ij}(t)] & i=2,\ldots,k-1 \\ m_{ij}(t)[g_{(i-1)j}(t)\lambda_{(i-1)j}(t) + \lambda_{ij}(t)] & i=k \end{cases} \tag{S1}$$

where $g_{ij}(t)$ represents the proportion of individuals in size class $i$ that move into size class $i+1$, $m_{ij}(t)$ represents the proportion of individuals in size class $i$ that survive, and $f_{ij}(t)$ represents the per capita regeneration rate of size class $i$ (i.e., the number of individuals it contributes to the smallest size class) over time step $t \to t+1$. Note that with the above definition of growth, $1 - g_{ij}(t)$ represents the proportion of individuals in size class $i$ that do not grow into the next larger size class ($i \to i$). Further, $g_{kj}(t)$ is equal to zero since individuals cannot grow beyond the largest size class $k$.

The above set of equations (Eqtn. S1) highlights the sparsity of the propagator matrix as defined in Eqtn. 5. Specifically, the growth matrix, $\mathbf{G}_j(t;\boldsymbol{\theta}_j^{\mathrm{gr}})$, is lower bidiagonal (non-zero elements along the diagonal and subdiagonal only) with diagonal elements equal to $1-g_{ij}(t)$ for ($i=1,\ldots,k$) and subdiagonal elements equal to $g_{ij}(t)$ for ($i=1,\ldots,k-1$). The mortality matrix, $\mathbf{M}_j(t;\boldsymbol{\theta}_j^{\mathrm{mr}})$, is diagonal (non-zero elements along its diagonal only) with the diagonal elements equal to $m_{ij}(t)$ for ($i=1,\ldots,k$). Finally, the fecundity matrix, $\mathbf{F}_j(t;\boldsymbol{\theta}_j^{\mathrm{fr}})$, contains non-zero elements only in its first row with values equal to $f_{ij}(t)$ for ($i=1,\ldots,k$). In total, the propagator matrix consists of $(3\times k)-1$ demographic rate terms.

The process model can be compactly defined for the full size-species distribution (across all species) as,

$$\boldsymbol{\lambda}(t+1) = (\mathbf{H}(t;\boldsymbol{\theta})\boldsymbol{\lambda}(t)) \odot \boldsymbol{\eta}(t+1) \tag{S2}$$

where $\mathbf{H}(t;\boldsymbol{\theta})$ is an $(mk\times mk)$ dimensional block diagonal matrix with species-specific propagator matrices along the diagonal,

$$\mathbf{H}(t;\boldsymbol{\theta}) = \mathrm{bdiag}(\mathbf{H}_1(t;\boldsymbol{\theta}_1),\ldots,\mathbf{H}_m(t;\boldsymbol{\theta}_m)),$$

$\boldsymbol{\theta}$ is a $mp$-dimensional vector including all species-specific demographic parameters, $\boldsymbol{\eta}(t+1)$ is an $mk$-dimensional vector including all species-by-size process error terms, and $\odot$ indicates the Hadamard (elementwise) product. The block diagonal structure of the full propagator matrix underscores both the sparsity of the propagator matrix and the fact that species dependence is modeled through the demographic functions used to populate the matrix rather than through the matrix itself.

### S4 Demographic functions

#### S4.1 Initial, theoretical functions

We initially defined demographic functions inspired by the SORTIE model (Pacala et al., 1996), defined applying the perfect plasticity approximation (PPA) (Strigul et al., 2008). Under these initial functions, growth rate was modeled as,

$$g_{ij}(t) = \text{pgr}_{ij}\Phi(\theta^{\text{gr}}_{j1} + \theta^{\text{gr}}_{j2}\text{r2pca}_{ij}(t))$$

where $\Phi$ is the cumulative distribution function of a standard normal distribution, $\text{pgr}_{ij}$ is a size- and species-specific maximum potential growth rate, $\text{r2pca}_{ij}(t)$ is the ratio of the realized to potential exposed crown area of individuals in a given size class (estimated conditional on the theoretical canopy base height), $\theta^{\text{gr}}_{j1}$ describes the growth of understory trees (with no exposed crown area), and $\theta^{\text{gr}}_{j2}$ describes how growth rate scales with exposed crown area. We relied on previously published studies for potential growth rates (Bragg, 2001), and crown allometries (Purves et al., 2007). Survivorship was then modeled as a function of the modeled growth rate,

$$m_{ij}(t) = \frac{\theta^{\text{mr}}_{j1}}{1 + \exp(-(\theta^{\text{mr}}_{j2} + \theta^{\text{mr}}_{j3} g_{ij}(t)))}$$

where $\theta^{\text{mr}}_{j1}$ describes the maximum survivorship rate, $\theta^{\text{mr}}_{j2}$ describes the survivorship of size classes exhibiting no growth, and $\theta^{\text{mr}}_{j3}$ describes how survivorship scales with growth rate. Lastly, fecundity was modeled as an exponential function of size,

$$f_{ij}(t) = \exp(\theta^{\text{fr}}_{j1} + \theta^{\text{fr}}_{j2} x_i)$$

where $x_i$ is the standardized DBH of the $i$th size class, $\theta^{\text{fr}}_{j1}$ is an intercept term, and $\theta^{\text{fr}}_{j2}$ describes how fecundity scales with DBH. Note that while the per capita fecundity rate within a size class is constant with respect to time, temporal variability in regeneration arises through the latent density of individuals in different size classes ($\boldsymbol{\lambda}_j(t)$). While recent work indicates that fecundity rates decline as trees near their maximum diameter (Qiu et al., 2021), we excluded higher order size terms in the name of parsimony given that the diameters of trees in the modeled forests were much smaller than their maximum.

We found that the above functions applying the PPA as a measure of density-dependence led to identifiability issues in demographic parameters when applied to both the Penobscot Experimental Forest (PEF) and simulated datasets, the latter of which applied the above functions as the true data generating mechanism. Several factors may have contributed to the observed identifiability issues. There was limited variability in the crown area and resulting growth rates of modeled species with individuals of a given species present in either the overstory or understory. The lack of variability in crown area tended to increase correlation among modeled demographic parameters, which is a known challenge for the Hamiltonian Monte Carlo algorithm utilized to fit the dynamical model. Other MCMC algorithms to update the demographic parameters, such as a block Metropolis or elliptical slice sampler, may be better able to identify the demographic parameters under the PPA.

##### S4.2 Functions for simulated data

Given the above identifiability issues, we replaced the PPA with basal area larger as a measure of density-dependence (similar to Kohyama, 1993), and simplified the demographic functions. Under the simplified functions growth is modeled as,

$$g_{ij}(t) = \frac{\mathrm{pgr}_{ij}}{1 + \exp(-(\theta^{\mathrm{gr}}_{j1} + \theta^{\mathrm{gr}}_{j2}\mathrm{bal}_{ij}(t)))} \tag{S3}$$

where $\mathrm{bal}_{ij}(t)$ is the total basal area of all trees larger than the target size class (summed across all species), and $\theta^{\mathrm{gr}}_{j1}$ and $\theta^{\mathrm{gr}}_{j2}$ are species-specific growth rate parameters. Survivorship is modeled as a function of DBH,

$$m_{ij}(t) = \frac{1}{1 + \exp(-(\theta^{\mathrm{mr}}_{j1} + \theta^{\mathrm{mr}}_{j2}x_i + \theta^{\mathrm{mr}}_{j3}x_i^2))} \tag{S4}$$

where $x_i$ is the standardized midpoint of the $i$th size class, and $\theta^{\mathrm{mr}}_{j1}$, $\theta^{\mathrm{mr}}_{j2}$, and $\theta^{\mathrm{mr}}_{j3}$ are species-specific mortality parameters. Lastly, fecundity is modeled as a species-specific, size-invariant per capita rate for size classes above a minimum value,

$$f_{ij}(t) = \begin{cases} 0 & d_i < d_{\min} \\ \exp(\theta^{\mathrm{fr}}_{j1}) & d_i \geqslant d_{\min} \end{cases} \tag{S5}$$

where $d_i$ is midpoint of the $i$th size class, $d_{\min}$ is the minimum size for regeneration (25.4 cm in our analysis), and $\theta^{\mathrm{fr}}_{j1}$ is a species-specific fecundity rate parameter.

##### S4.3 Functions for Penobscot Experimental Forest

Applying the dynamical model to PEF data requires several simplifications to the above-defined demographic functions. First, there was limited observed regeneration in the management unit over the model period. Specifically, ingrowth into the smallest DBH class was observed in just three inventory years (1984, 1993, 1999) for two shade-tolerant species (balsam fir, eastern hemlock) in a maximum of two sample plots. Given limited observations to inform the regeneration component of the process model and the pulsed nature of regeneration events, we turn this model component off when applied to the PEF data.

Initial applications of the dynamical model to the PEF data indicated insufficient information to inform the defined growth parameters (Eqtn. S3). The lack of identifiability appears to be related to limited variability in the size of all species except for eastern white pine, which forms the main canopy of the even-aged stand (Fig. S2). Further, the individual potential growth rate term (pgr in Eqtn. S3) appears overly restrictive and not closely connected to the population-level growth modeled here. As such, we drop the potential growth rate and basal area larger terms from the function such that growth is modeled as a species-level mean growth rate $\theta^{\text{gr}}_{j1}$. We allow size to inform white pine growth rates by including DBH class as an additional predictor,

$$g_i(t) = \Phi(\theta^{\text{gr}}_1 + \theta^{\text{gr}}_2 x_i(t)),$$

given that there was sufficient variability in white pine diameters to estimate the second growth parameter.

Lastly, initial applications of the dynamical model indicated that mortality rates did not vary strongly as a function of size over the model period. As such, we drop the size terms in the mortality function (Eqtn. S4) modeling survivorship as a species-level mean over the model period.

We again attribute identifiability issues to the lack of variability in the data. Specifically, the lack of basal area and size variability among species over the model period. The lack of variability in the growing conditions within the PEF stand points to the need to integrate information from a network of plots providing observations of forest ecosystems in a diversity of states.

#### S5 Model implementation

##### S5.1 Prior distributions

We apply a non-centered parameterization to estimate demographic parameters for each species conditional on a shared mean $\mu_\ell$ and standard deviation $\tau_l$,

$$\theta_{j\ell} = \mu_l + z^\theta_{j\ell}\tau_l,$$

where $z^\theta_{j\ell}$ is assigned an independent standard normal prior and $\ell$ is used to index the demographic parameters ($\ell = 1, \dots, p$). The parameter scaling white pine growth by DBH ($\theta^{\text{gr}}_2$), as defined above, is assigned a semi-informative normal prior: $\theta^{\text{gr}}_2 \sim \text{N}(0.5, 10)$. Demographic parameters are unitless describing proportional growth and mortality on a probit and logit scale, respectively.

Process and observation error standard deviations are both assigned zero-truncated normal distributions,

$$\phi_j \sim \text{N}_{[0,\infty]}(0, 0.5)$$
$$\sigma_j \sim \text{N}_{[0,\infty]}(0, 0.5)$$

for each species. The observation and process error standard deviations are unitless; the former because it describes variation of a multiplicative observation error term, which is a ratio of the mean, and the latter because it represents the standard deviation of the latent density on the log scale. The process error terms are sampled using a non-centered parameterization,

$$\eta_{ij}(t) = \exp(z^{\eta}_{ij}(t)\,\sigma_j)$$

where $z^{\eta}_{ij}(t)$ is assigned an independent standard normal prior. The observation error terms are integrated out as described in the main text and are not estimated as part of the MCMC sampler.

Initial condition parameters include the total stem density for each species at time zero ($n_j$) expressed in stems per hectare, and the mean ($q_j$) and standard deviation ($v_j$) of the standard normal distribution used to approximate the initial size density (Eqtns. 11 and 12), both of which are unitless since they are associated with a standard normal reference distribution. We use a non-centered parameterization to estimate the initial stem density and mean of the size distribution,

$$\begin{aligned} n_j &= \exp(\tilde{n}_j + z^n_j\,\tau^n_j) \\ q_j &= \tilde{q}_j + z^q_j\,\tau^q_j \end{aligned}$$

where $\tilde{n}_j$ and $\tilde{q}_j$ are prior estimates of the initial values and $\tau^n_j$ and $\tau^q_j$ are prior uncertainty estimates based on the first inventory observation as described in the main text. Note that the mean of the standard normal distribution used to approximate the initial size distribution may be negative as is likely to be the case for strongly right-skewed size distributions. We assign $z^n_j$ and $z^q_j$ independent standard normal priors for each species. Lastly, we assign the a half Cauchy prior to the standard deviation of the initial size distribution: $v_j \sim \text{HalfCauchy}(0, 5)$.

##### S5.2 Starting values

Starting values for demographic rate parameters are generated based on generalized linear growth and mortality models fit to individual tree demographic estimates based on PEF inventory observations. Starting values for process and observation error components as well as the standard deviation of the initial size distribution are sampled from assigned prior distributions. Lastly, starting values for the initial value parameters are generated by fitting the initial size distribution model (as defined in Eqtns. 11 and 12) to the first inventory observation. Starting values for the simulated data analysis were generated by adding random noise to simulation parameter values.

##### S5.3 Recovering observation errors

As described in the main text, the Poisson data model with a Gamma prior assigned to the multiplicative observation error leads to a negative binomial model with the observation errors marginalized out. These observation errors can be recovered via compositional sampling following model convergence. Specifically, the observation errors follow a gamma distribution conditional on the overdispersion parameter ($\phi_j^2$) and inventory observations ($y_{ij}^{\ell}(t)$:

$$\epsilon_{ij}^{\ell}(t)|\phi_j^2, y_{ij}^{\ell}(t) \sim \text{Gamma}(\alpha, \beta)$$

$$\alpha = \frac{1}{\phi_j^2} + y_{ij}^{\ell}$$

$$\beta = \frac{1}{\phi_j^2} + |A|\,\lambda_{ij}(t).$$

We apply posterior samples of observation error terms to construct plot-level predictions of stem and basal area density (Figs. S7, S8).
